# What comes after *de novo*? Automated lead optimization of proteins with CRADLE-1

**DOI:** 10.64898/2026.03.06.710001

**Authors:** Eli Bixby, Gino Brunner, Daniel Danciu, Richard Dela Rosa, Nicolas Deutschmann, Constance Ferragu, Franziska Geiger, Christian Holberg, Patrick Kidger, Arthur Lindoulsi, Noé Lutz, Thomas McColgan, Sebastian Millius, Harmen van Rossum, Jinel Shah, Michelle Vandeloo, Paula Vidas, Daan van der Vorm, Jonathan D. Ziegler, Nicolò Baldi, Catalina IJSpeert, Emanuele Monza, Angela Schriek

## Abstract

Lead optimization remains the longest and most expensive step in pre-clinical drug discovery, typically consuming 12-36 months whilst costing $5M-$15M per candidate. We introduce ‘CRADLE-1’, an automated framework for protein engineering. While CRADLE-1 supports the full process of drug discovery and industrial protein engineering pipelines, including hit identification and *de nouo* binder design, this work focuses on its application to multi-property lead optimization across protein modalities (VHHs, scFvs, IgGs, peptides, enzymes, CRISPR systems, vaccines). We show it is 4-7× faster than rational design, as measured by the number of wet lab rounds required. We provide in-vitro validation across all of the above modalities, typically optimizing multiple properties simultaneously (single and polyspecific binding down to picomolar, activity, thermostability,.). Technically, CRADLE-1 starts with pre-trained foundation protein language models (PLMs), which are fine-tuned in unsupervised fashion on evolutionary neighborhoods, in supervised fashion using lab-in-the-loop data, and then deployed in a multi-model workflow. Of additional interest, we find that (a) the end-to-end system may be run in automated fashion; (b) wet lab data may be consumed in ‘black box’ fashion without knowledge of the underlying biochemical mechanisms; (c) structural data may largely be superseded by sequence-function pairs.

## 1 Main

Lead optimization is the iterative refinement of a candidate molecule - such as an antibody, enzyme, or vaccine - to enhance its functional profile. While a “lead” identifies a promising functional starting point, it rarely possesses the properties required for practical application. Lead optimization involves the simultaneous tuning of multiple traits: for a therapeutic antibody, this may mean increasing affinity and developability, while reducing immunogenicity; for an industrial enzyme, it may involve increasing catalytic turnover while maintaining stability in harsh pH or temperature conditions. Paul, S. M. *et al*. 1 estimate that lead optimization is the longest and most expensive step of the pre-trial drug discovery pipeline, requiring 12–36 months of iteration, at a cost of $5M–$15M per candidate, and with 1 out of 14.6 candidates leading to a successful product launch. Lead optimization is thus a major cost driver for drug development.

The lead optimization process itself is typically pursued through designing variants of a candidate molecule, building and testing these candidates in the lab, and then using the gathered information to design a next round of variants. This process is often referred to as design-build-test-learn cycles (Figure 1).

**Figure 1:**
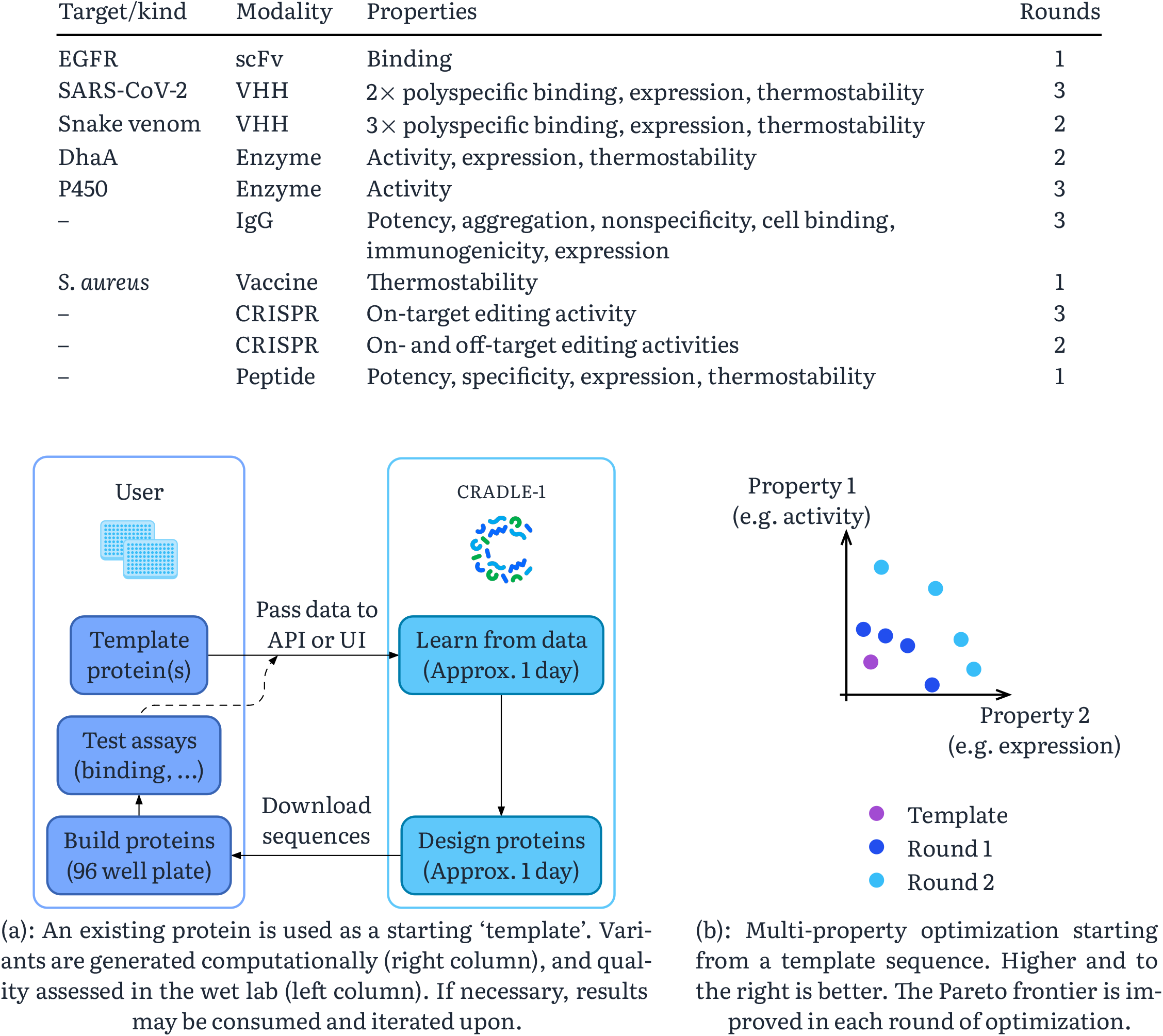
The lead optimization (design-build-test-learn) process, as typified by the interaction between CRADLE-1 and a user.

### 1.1 Contributions

Machine learning – here, the CRADLE-1 system – may be used to design candidate proteins and to learn from wet lab results. When doing so, we provide evidence in support of two findings:

1. The lead optimization process is faster (4–7x, as quantified by number of wet lab rounds) than typical alternatives, such as human-in-the-loop rational design, even when supported through ML/computational tools such as structure prediction or variant effect prediction. Moreover, this is true across:
  - multiple modalities (VHHs, scFvs, IgGs, peptides, enzymes, CRISPR systems, vaccines);
  - simultaneous optimization of multiple properties (1-6 discussed here; successfully up to 8 in private benchmarks);
  - varying numbers of consecutive wet lab rounds (1-3, with more having never been required);
  - high numbers of residues mutated per round (1-15 discussed here; successfully up to 52 in private benchmarks);
  - the availability or non-availability of in-context sequence-function data;
2. These computational aspects are automatable, and that this is true even in the presence of noisy wet lab data exhibiting batch effects. Practically speaking, this is an API call or UI interaction which consumes wet lab data and which returns designed proteins.

We hypothesize that these improvements are possible as lead optimization is traditionally heuristic (centered on human-in-the-loop design, using computational tools), and machine learning is excellent at heuristics.

Our analysis encompasses a diverse array of protein modalities, extending beyond therapeutic candidates to include industrial enzymes and gene-editing systems. However, we only consider proteins, and do not consider small molecules. Here we focus on lead optimization; other aspects of CRADLE-1, such as hit identification or *de nouo* binder design will be reported elsewhere. Results are provided in Section 2. Algorithmic and machine learning details are summarized in Section 3. The following table provides a summary of results; ‘-’ indicates a redaction on behalf of a commercial partner.

### 1.2 Lead optimization and *de novo* binder design

*De nouo* binder design has been the focus of much of the literature, with examples including machine-learning-guided design of peptides^2–4^, VHHs, scFvs^5–8^ and mAbs^9^. Such approaches are often enabled via underlying foundation models for structure prediction^10–12^, docking^13^, backbone prediction^14^, or inverse folding^15^.

For clarity, we emphasize that the present work tackles a different problem. Our focus is more broad (optimizing any kind of protein modalities with respect to any kind of property, rather than antibody fragments for binding) and at a different step in the drug discovery pipeline (lead optimization, rather than hit discovery). There is comparatively less literature on this topic^16–20^. The technical methods are also different (Section 3), with a focus on protein language models21 instead of structure-based methods.

## 2 Results

CRADLE-1 has successfully designed dozens of campaigns for commercial and academic partners, in addition to numerous internal studies. In this paper, we present a representative subset of these results, selected from in-house experiments and specific commercial cases (redacted where appropriate) with partner consent for disclosure.

### 2.1 scFv framework optimization for binding to a transmembrane protein

We engineer 12 variants of a single-chain variable fragment (scFv) to bind to epidermal growth factor receptor (EGFR), which is a transmembrane protein whose overexpression is associated with numerous cancers.^22^ All 12 variants achieve binding, with values ranging from 339 pM to 4.51 nM. This was performed as part of the Adaptyv Bio protein design competition^23^; there were 400 entries, which were required to differ from any known therapeutic binder by a Levenshtein distance of at least ten. CRADLE-1 won the competition; the second-place entry had a binding of 5.18 nM.

EGFR has a known commercial binder, Cetuximab (sold under Erbitux), which we format as an scFv with a glycine-serine linker, and use as the template sequence. This exhibits binding of 6.64 nM. Variants are obtained whilst holding the complementary-determining regions (CDRs) of the scFv constant, and optimizing only the remaining (‘framework’) regions. We hypothesized that whilst the CDRs capture gross binding behavior, framework mutations may then allow for more subtle structural changes that could fine-tune binding. All binders are ten mutations away from the template sequence, and up to 19 mutations away from each other. In total, 33 unique mutations across 25 distinct positions were generated.

See Figure 2. Further results may be found in Appendix B.1, including an *in silico* evaluation of immunogenicity.

**Figure 2:**
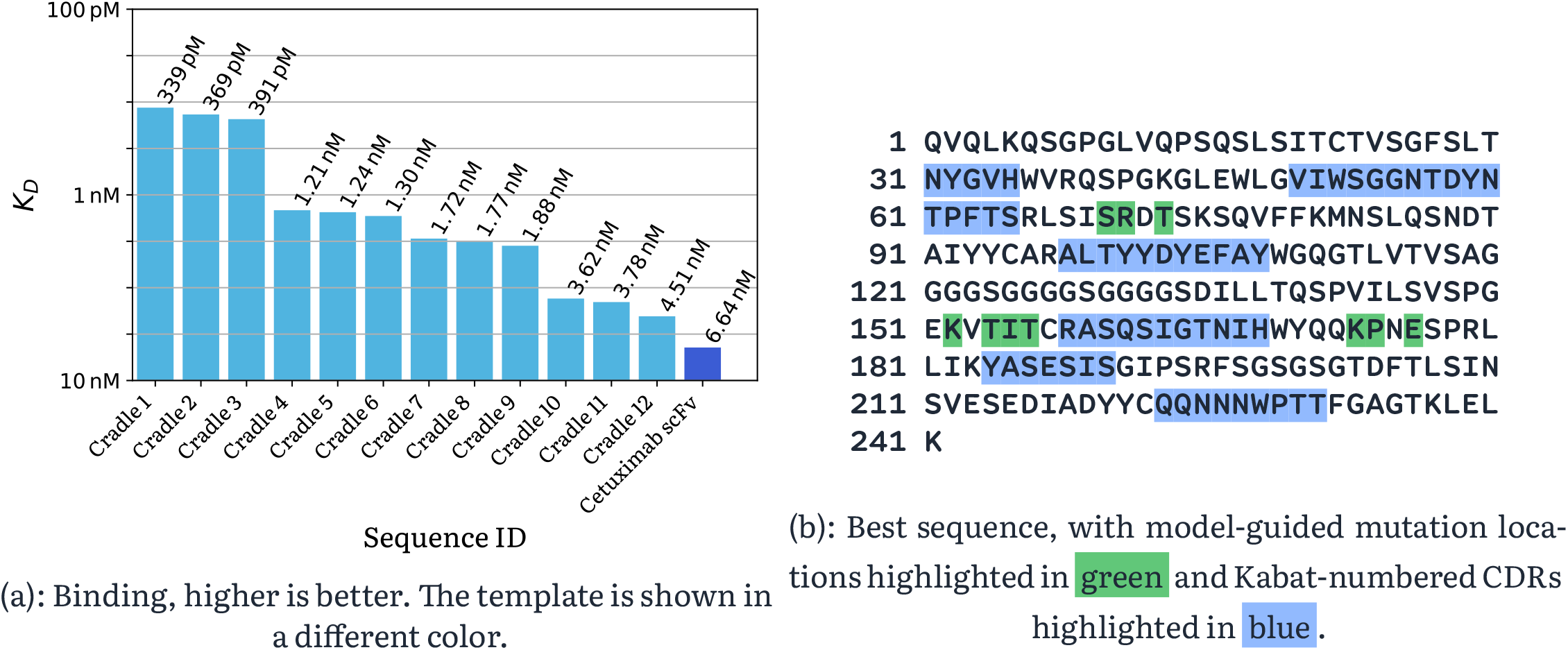
cradle-1-generated scFvs, and template scFv-formatted Cetuximab, for binding against EGFR.

### 2.2 VHHs with picomolar and polyspecific binding

#### 2.2.1 SARS-CoV-2

We engineer a VHH (‘nanobody’), simultaneously optimizing four properties: binding to SARS-CoV-2 wild type, binding to SARS-CoV-2 Omicron, thermostability, and expression in *E. coli*. We obtain a VHH exhibiting binding against wild type with a *K*_*D*_ of 186 pM, binding against Omicron with a *K*_*D*_ of 11.4 nM, a melting temperature of 70.9 °C, and 1.88x improved expression.

This is intended as a challenging problem. The template sequence was derived from phage display NGS data using CRADLE-1’s hit identification module (not discussed here). This sequence already exhibits good (but not excellent) properties, so that ‘most’ mutations are assumed to be deleterious to one or more properties, with relatively few good options for improvement. Our template sequence was obtained from Hanke, L. *et al*. ^24^, and exhibits binding against SARS-CoV-2 wild type with a *K*_*D*_ of 2.01 nM, binding against Omicron with a *K*_*D*_ of 34.6 nM, and a melting temperature of 58.5 °C.

We plot binding to SARS-CoV-2 wild type against thermostability in Figure 3. Three rounds of iteration progressively push out the Pareto frontier. The first round uses only evolutionary context as data; the latter two rounds fine-tune models on the gathered wet lab data. The final best sequence is chosen by manually selecting a sequence from the Pareto frontier, with a reasonable trade-off amongst all properties.

**Figure 3:**
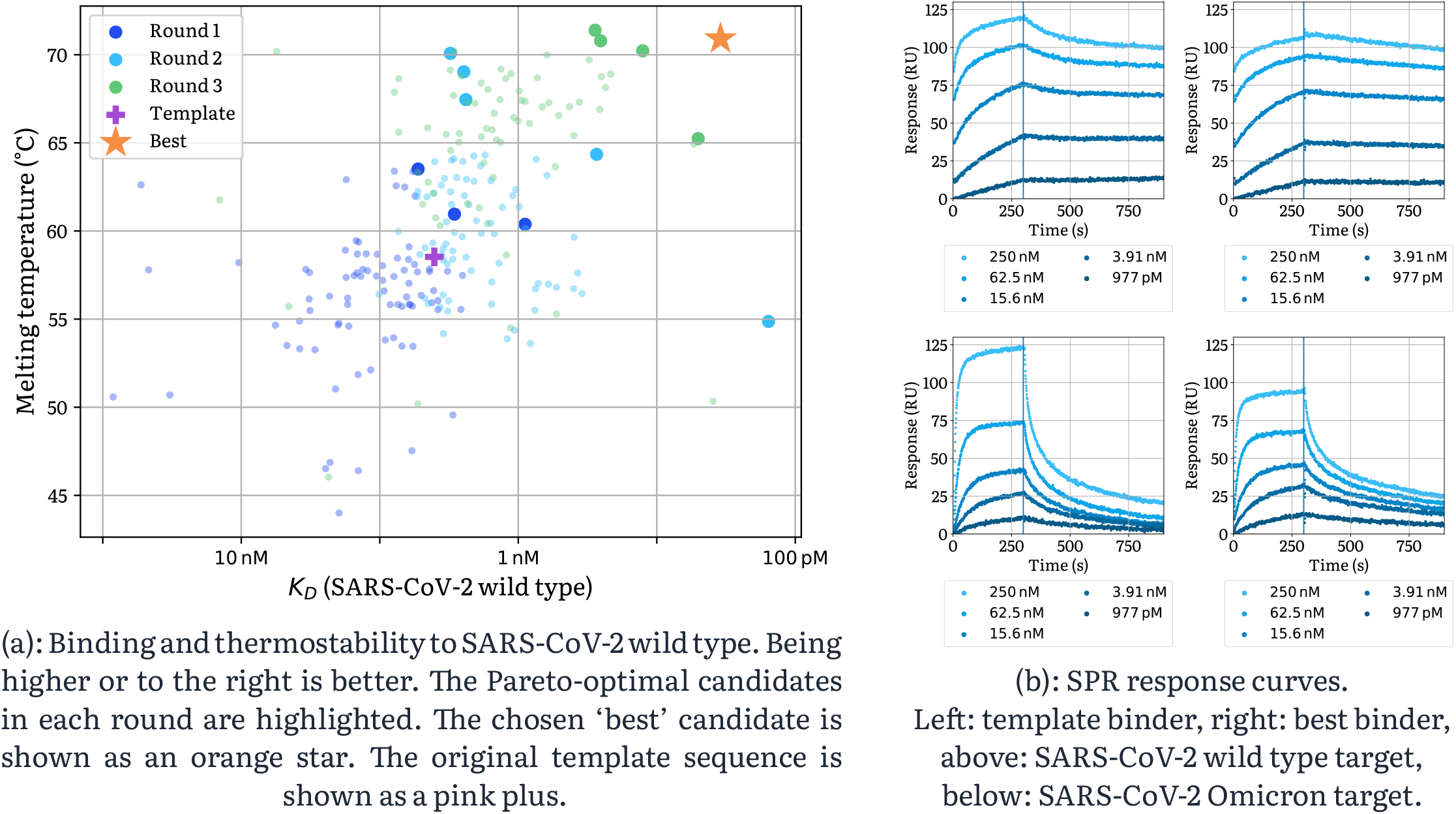
VHHs obtained with cradle-1 for polyspecific binding to SARS-CoV-2 wild type and SARS-CoV-2 Omicron.

Further details are available in Appendix B.2.1, including an all-pairs Pareto plot. A comparison against computational and protein engineering baselines is available in Appendix A.1.

#### 2.2.2 Snake venom

Snakebites are estimated to kill 7 000 people every year in sub-Saharan Africa^25^ and 58 000 people every year in India26. The species of snake may not always be known when administering antivenom for treatment, and for this reason polyvalent or polyspecific antivenoms are an important therapeutic modality.^27^ High thermostability is also of importance, as cold storage facilities may not be available.

We perform a five-property optimization for VHHs with binding against three different long-chain three-finger *α*:-neurotoxins (LNTxs) which are common among elapids, along with thermostability and expression. The template sequences were again obtained using CRADLE-1’s hit identification module applied to NGS data obtained from a phage display campaign (not discussed here). Mutations are made to both the CDR and framework regions. Three template sequences are initially obtained via a phage and yeast display campaign. We obtain a VHH with <100 pM, <1 nM, <2 nM bindings against each of the three toxins, a melting temperature of 76.7 °C, and 1.59x fold improvement in expression relative to the templates.

This optimization campaign was run in conjunction with an academic partner, and full details will be released in an upcoming publication. Auxiliary plots may be found in Appendix B.2.2.

### 2.3 Joint optimization of thermostability and expression of a haloalkane dehalogenase

We optimize the thermostability and expression of a haloalkane dehalogenase (accession Q53042), whilst preserving its activity.

The template sequence has a melting temperature of 45.1 °C. Two rounds were generated using CRADLE-1, and an improved sequence was obtained. The melting temperature was improved by +20.0 °C to 65.1 °C, expression was improved by a factor of 2.06x, and activity was improved by a factor of 1.29x.

Amongst the CRADLE-1-generated sequences with improved expression over the template, the candidates with the highest 39 thermostabilities were selected for an activity assay. See Figure 4, and further details in Appendix B.4.

**Figure 4:**
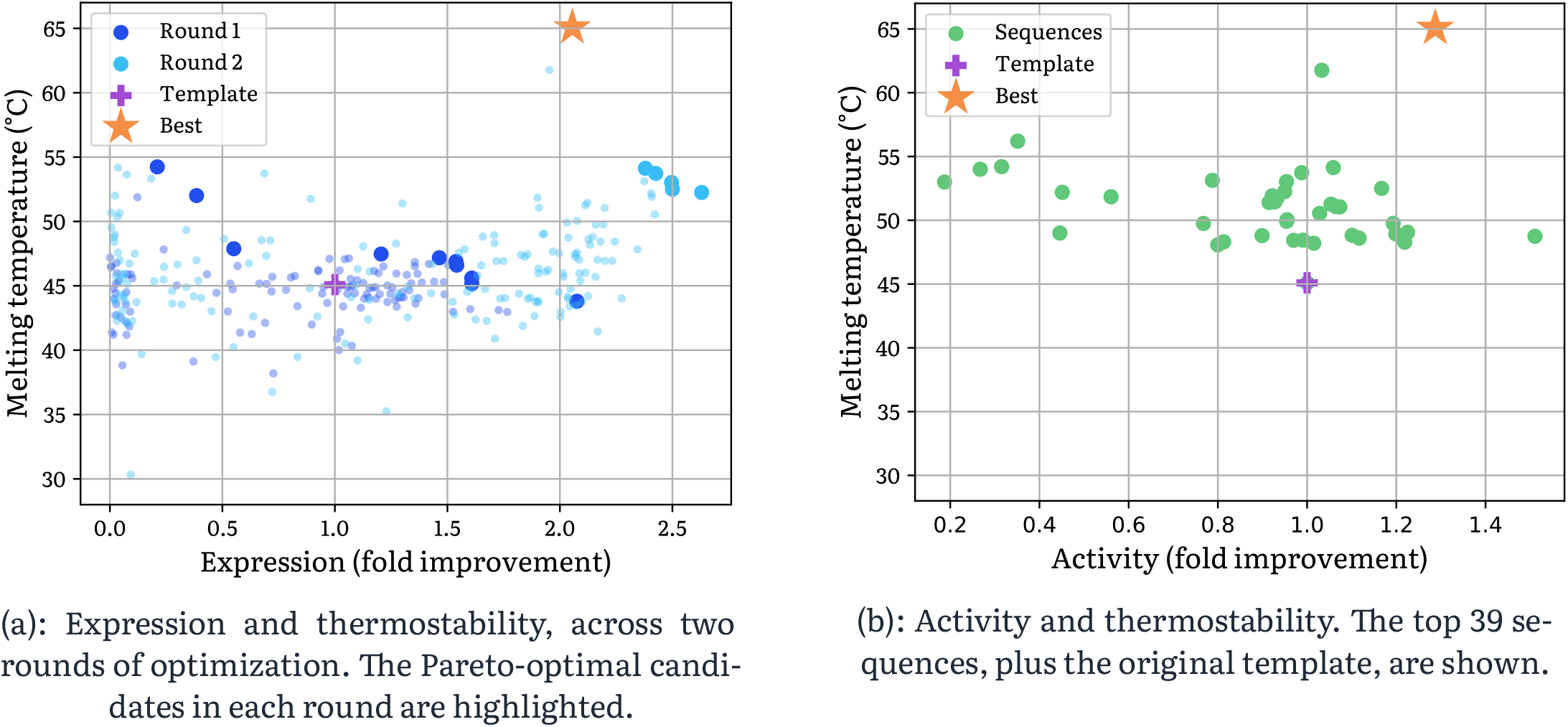
cradle-1-generated haloalkane dehalogenases are optimized across two rounds of optimization. Being higher or to the right is better. The chosen ‘best’ candidate is shown as an orange star. The original template sequence is shown as a pink plus.

### 2.4 Optimizing activity of a P450 enzyme

We performed single-property optimization of the activity of a P450 enzyme on behalf of a commercial partner. The partner had previously used rational design, trialing 1201 candidates over eight experimental rounds, obtaining a maximal fold improvement of 17.9x. This was considered a large number of rounds, and the fold improvement considered insufficient, and for this reason the project was originally scheduled for cancellation.

Three rounds were then generated using CRADLE-1, with 96 candidates each. This achieved a fold improvement of 40.6x. The median round-over-round increase using CRADLE-1 was 3.82x higher than in the rational design effort.

See Figure 5. Further details are available in Appendix B.5.

**Figure 5:**
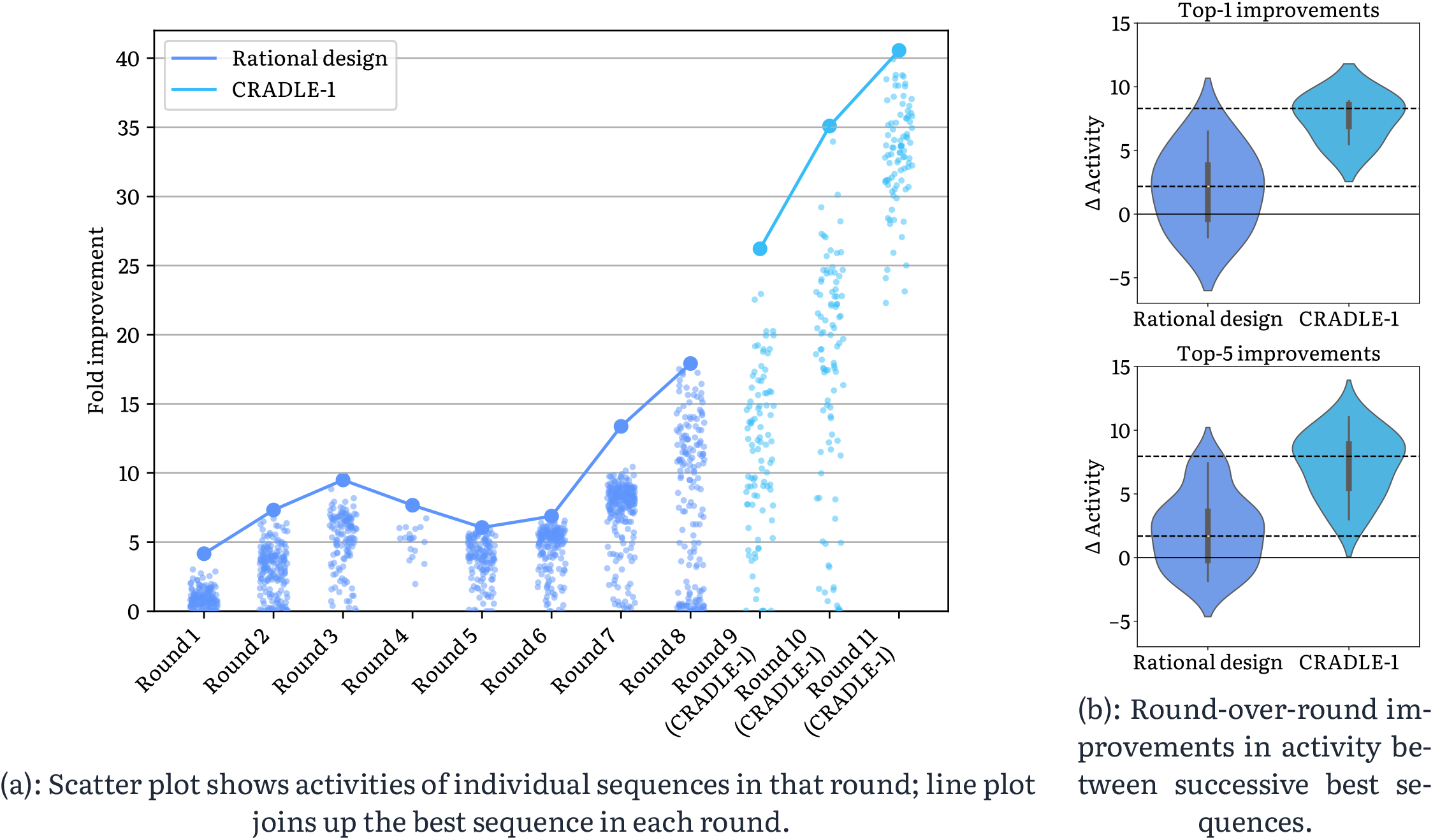
Activity of P450 variants. Initial rounds are rationally-designed sequences, followed by cradle-1-generated sequences. Higher is better. The problem is assumed to be challenging; rounds 4-6 get worse due to the rationally-designed sequences failing to produce any hits.

### 2.5 Aggregation, nonspecificity, immunogenicity, and potency of an lgG

Using CRADLE-1, we successfully engineered an IgG for potency (tuned to a range of one order of magnitude), aggregation (SMAC and AC-SINS), polyreactivity/nonspecificity (PAIA), cell binding, immunogenicity, and expression, on behalf of a top-50 pharmaceutical commercial partner.

A single functional parental sequence was available and was used as the template sequence. This variant exhibited unfavorable manufacturability characteristics, including two existing sequence liabilities which ideally should be engineered out.

The partner had previously run a screening campaign, which had failed to obtain variants demonstrating potency (even with successful binding as confirmed via ELISA). The partner had also attempted in-house lead optimization, and this similarly failed to obtain satisfactory variants.

Three rounds were generated using CRADLE-1 with 96 candidates each. In the end, 10 successful candidates meeting both potency and developability criteria were obtained and brought forward to the next stage of the pipeline.

### 2.6 Multi-property optimization of a bispecific VHH

We have successfully engineered a bispecific VHH, optimizing for binding to two distinct targets, while maintaining or improving thermostability, expression, polyreactivity, and hydrophobicity. This was done without any existing data on the bivalent construct. We compared the performance of CRADLE-1 when supplied with no data beyond the template sequences, versus when supplied with screening data on the individual monospecific VHHs and variant data with characterized binding, thermostability, and expression for the individual monospecific VHHs. The results, presented in Figure 6, show that even without any data, CRADLE-1 is able to generate candidates with good binding to both targets, while preserving or improving additional properties as seen in Figure 17 in Appendix B.3. However, the inclusion of in-domain context data significantly improves performance, yielding candidates with more favorable trade-offs between all properties. This experiment demonstrates that CRADLE-1 is able to learn from labeled data and partial sequence information, providing a strong starting point for the design of more complex proteins.

**Figure 6:**
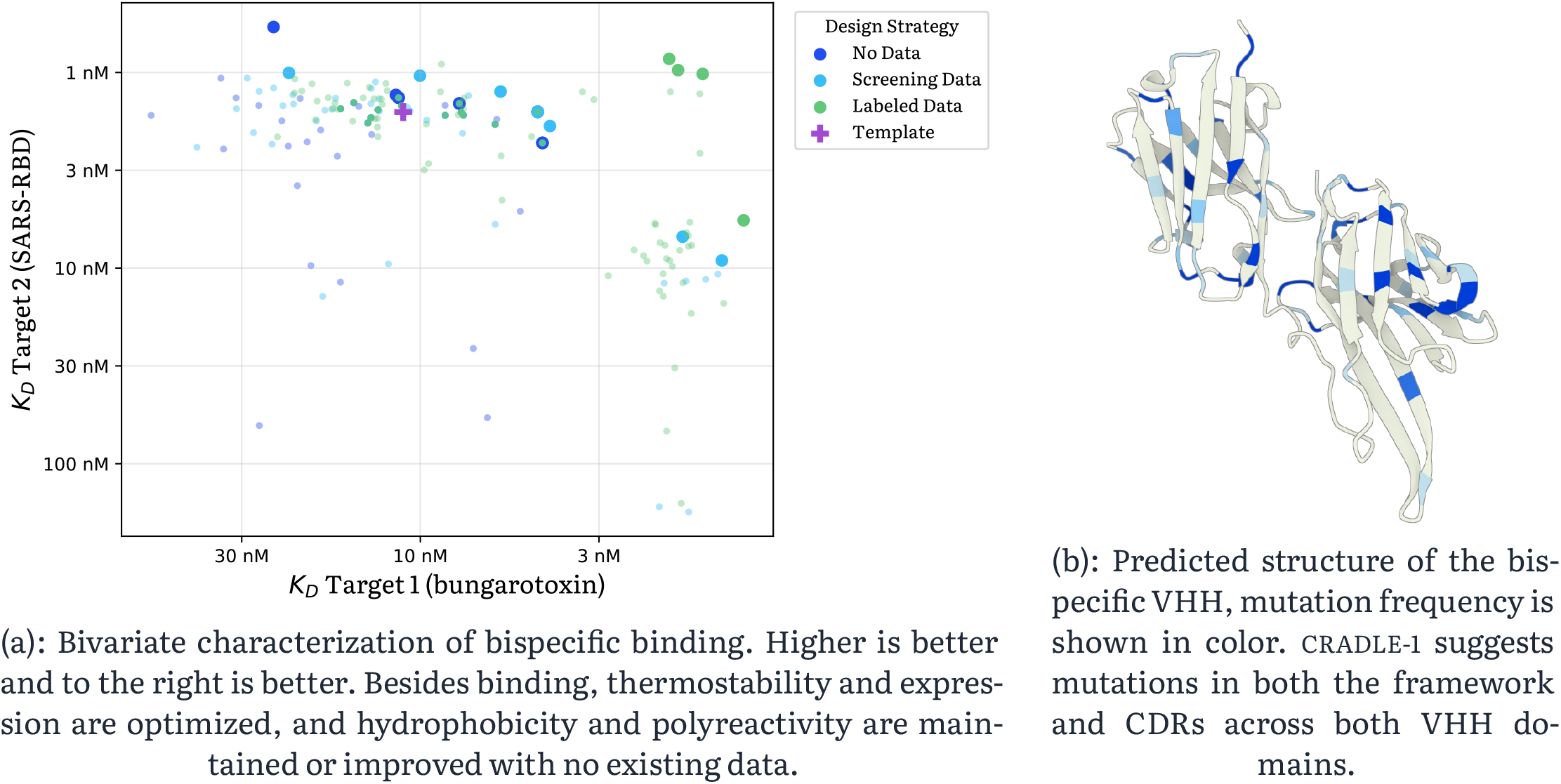
Bivariate evaluation of a bispecific VHH binding to two distinct targets. This demonstrates that cradle-1 is able to learn from partial data on single-domain variants to improve design of more complex bispecific variants.

### 2.7 Thermostability of a chimeric *Staphylococcus aureus* vaccine

The pathogenic bacterium *Staphylococcus aureus* (*S. aureus*) causes upper respiratory and gut infections and was associated with over 1 million deaths in 2019.

We have successfully optimized a chimeric vaccine candidate for *S. aureus* with a top-20 pharmaceutical commercial partner, improving the thermostability of an antigen by 2.50 °C in a single round of optimization. The partner estimated that this was 7x faster (as measured by number of rounds of wet-lab optimization) than a historical rational design effort.

### 2.8 Editing activity and specificity of a CRISPR system

We successfully engineered two CRISPR systems for on- and off-target editing activities, on behalf of a commercial partner.

For the first project, the focus was on single-property optimization of on-target editing activity. In a previous optimization campaign screening >3000 variants, the partner’s best variant had an on-target editing activity of <25%. The data from this campaign was provided to CRADLE-1, which was used to generate three rounds of 96 candidates each. The first round produced a variant exhibiting an on-target editing activity of 50%, and this was further improved to 56% and 68% in the second and third rounds respectively.

The second project required both an increase in on-target editing activity and a decrease in off-target editing activity, at five distinct sites. A previous optimization campaign (utilizing structural and rational design approaches) obtained a hit achieving on-target editing activity of 40% and worst-site off-target editing activity of 0.4%. The data from this campaign was provided to CRADLE-1, which was used to generate two rounds of 96 candidates each. The best CRADLE-1 design improved on-target editing activity to 75%, and reduced the worst-site off-target editing activity to 0.1%.

### 2.9 Peptide optimization under tight multi-property constraints

Using CRADLE-1, we supported three peptide optimization projects for a top-20 pharmaceutical commercial partner. All projects were late-stage and close to a dead end, following multiple prior rounds of optimization with limited remaining sequence flexibility.

The key challenge across projects was meeting tight, simultaneous constraints on potency, specificity, expression, and thermostability. Previous in-house and external optimization efforts had failed to deliver sequences satisfying the full specification.

CRADLE-1 generated candidate libraries with a 50% success rate in matching all constraints, whilst delivering significant potency improvements among the best constraint-compliant sequences. The partner estimated that CRADLE-1 enabled them to iterate 5x faster, as measured by number of rounds of wet lab optimization. Multiple optimized peptides were advanced to downstream validation.

### 2.10 Baselines

We compared CRADLE-1’s performance against a computational baseline and a typical protein engineering baseline. We optimized an anti-SARS-CoV-2 VHH and an anti-Her2 scFv for 3 different properties across 3 rounds. The optimized properties were: expression (in *E. coli*), melting temperature and binding affinity. For the anti-SARS-CoV-2 VHH we optimized for polyspecific binding against both the wild type and the Omicron variant. We used the open source ProteusAI^28^ with an ESM-2^21^ model as a computational baseline and alanine scanning (anti-Her2) or CDR3 scanning (anti-SARS-CoV-2) followed by 2 recombinant rounds as a protein engineering baseline. The baseline methods received a quota of 192 sequences each for Round 1 (a CDR3 scan requires 11 × 19 = 209 sequences), while CRADLE-1 only received a quota of 96 sequences. All methods received a quota of 96 sequences in Rounds 2 and 3. Figure 7 and Figure 9 show CRADLE-1’s performance relative to the baselines. Further details are available in Appendix A.1.

**Figure 7:**
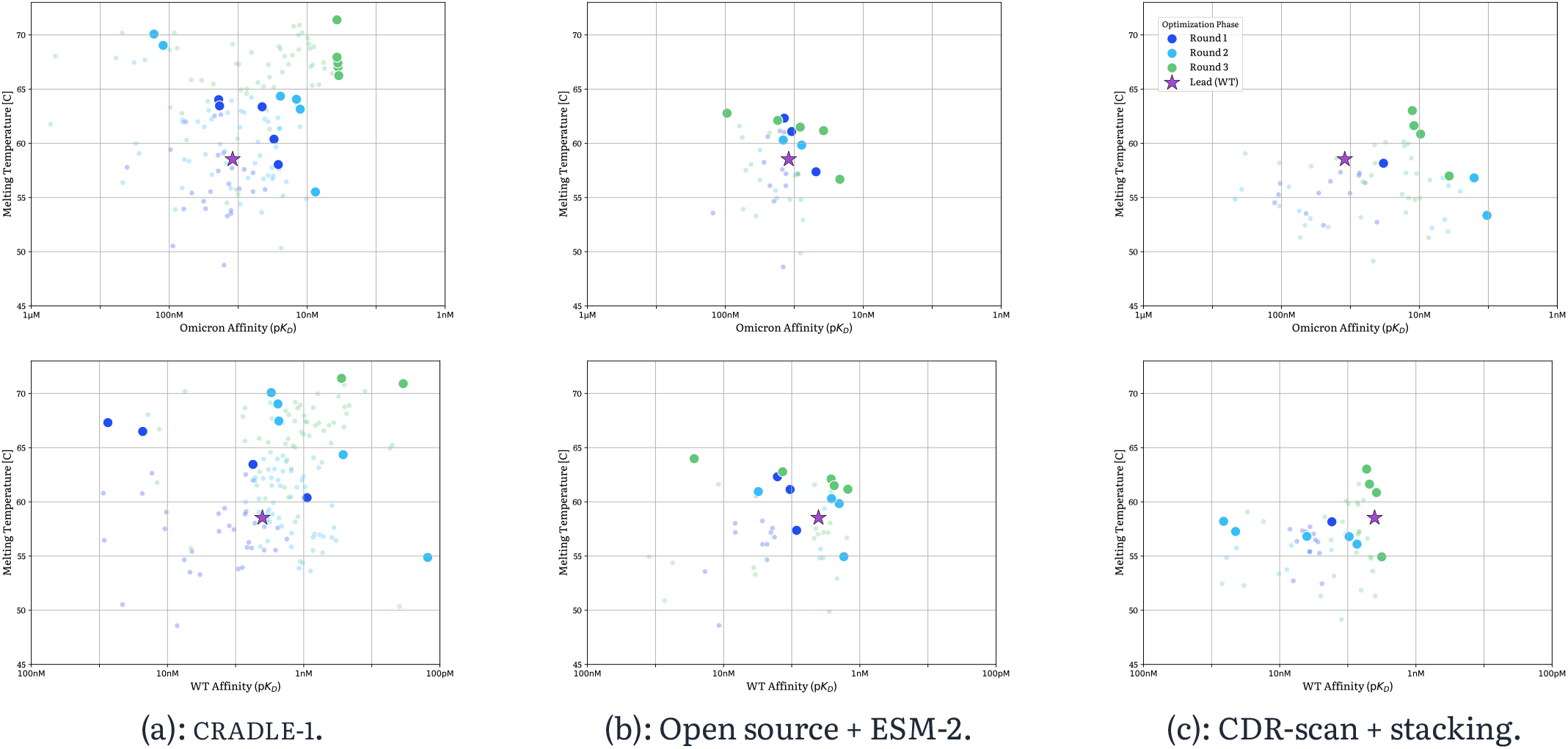
Comparative Pareto frontier analysis for multi-objective VHH optimization. Successive rounds of optimization for an anti-SARS-CoV-2 VHH are shown across three methodologies: cradle-1 (left), an open-source ESM-2-based method (middle), and CDR-DMS with stacking (right). Top row illustrates the trade-off between Omicron RBD affinity (*pK*_*D*_) and thermal stability (*T*_*m*_), bottom row depicts WT RBD affinity versus *T*_*m*_. Pareto-optimal candidates are highlighted by increased marker size and saturation; the starting template sequence is denoted by a star. Variants with expression levels below the template threshold are not shown.

## 3 Methods

CRADLE-1 combines multiple models into an overall system. Given a protein (referred to as the ‘template sequence’), and potentially a limited amount of paired sequence-function data, the CRADLE-1 system generates variants of that protein which are predicted to have improved function.

**Pre-training:** a foundational protein language model is trained on large-scale protein sequence databases such as UniRef^29^.

**Fine-tuning:** the foundation model is fine-tuned on the in-domain data to obtain two distinct models.

1. The foundation model is trained (‘evotuned’) in unsupervised fashion, optimizing a masked language loss, on the evolutionary context of the template sequence^30^.
2. If sequence-function data are available, then two distinct copies of the evotuned model are fine-tuned in supervised fashion in two different ways:
  2.1. via preference optimization, optimizing log-likelihood-ratio, over preference pairs using group direct preference optimization (g-DPO), which we introduce in Ferragu, C. *et al*. 31. The resulting model is referred to as a ‘logiter’. The g-DPO algorithm may be of independent interest as a computationally-efficient approximation to DPO.
  2.2. by adding a regression head, directly optimized to predict properties given the sequence. The resulting model is referred to as a ‘predictor’. Some of the algorithms designed here may be of independent interest, most notably an automated system for batch effect robustness, and a multi-property Spearman rank correlation (which we use for model selection).

**Generation:** an iterative sampling loop is used.

1. Candidates are first generated by modifying the template sequence. We perform a beam search over mutations, and select sets of mutation that are deemed favorable by the logiter.
2. Candidates are then ranked into a diverse set of sequences that are predicted to have high function by the predictor.
3. Iteration proceeds until enough candidates have been generated. This takes the form of a ‘double beam’ search over candidate sequences. As the temperature for generation increases over time, we maintain two beams in parallel, one containing currently-accepted sequences, and the other containing ‘backup’ sequences that may become suitable for acceptance at higher temperatures.

A diagram is provided in Figure 8.

**Figure 8:**
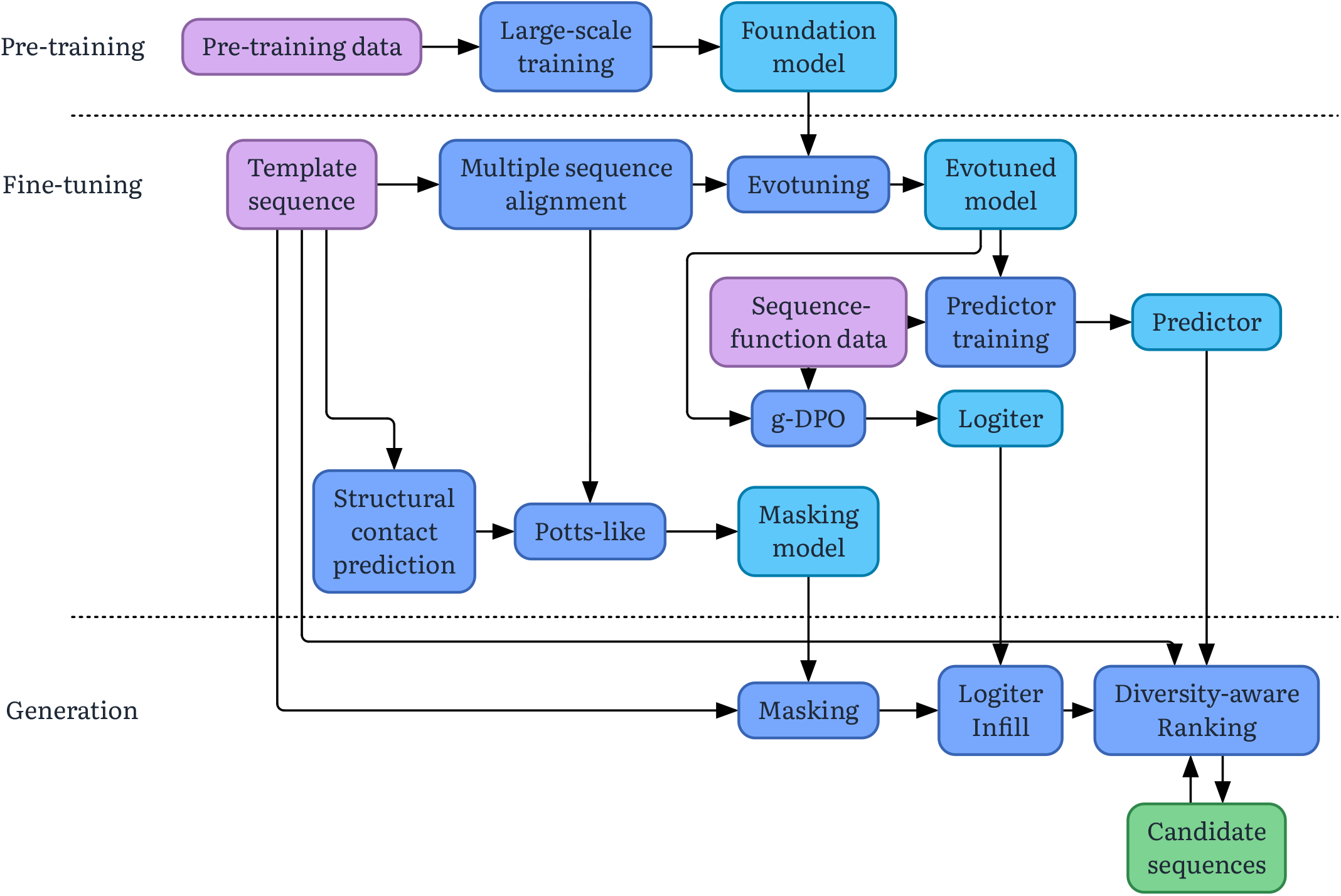
End-to-end system. Inputs to the system are displayed in pink, algorithmic processes in blue, models in sky, and outputs in green.

### 3.1 Zero-shot / diversification

The very first round of lead optimization will commonly have no data beyond the initial lead sequence(s). In particular, there may be no sequence-function data. When this is the case, the broad strokes of the CRADLE-1 system remain unchanged. g-DPO is skipped, and the evotuned model is directly used as a logiter. There is no predictor, and diversity-aware ranking is skipped.

For example, this is the setting in which CRADLE-1 won the Adaptyv challenge^23^, see Section 2.1. Similarly, it was the the first round of optimization in Section 2.2, Section 2.3, and Section 2.7.

We assume that the success of zero-shot depends on the extent to which the properties to optimize are evolutionarily conserved. Whilst properties such as thermostability are well-known to be evolutionarily conserved^32^, recent work has also shown that binding is conserved as well^16,33,34^.

## 4 Discussion

**Speed:** the end-to-end pipeline requires an aggregate compute effort of approximately two GPU-days (NVIDIA A100). However, through distributed parallelization across multiple GPUs, the actual wall-clock latency is reduced to the order of hours. This number is variable, primarily depending on (a) the amount of data, (b) the length of the template sequence, and (c) the hyperparameters of the CRADLE-1 system.

For the initial training round, we find that additional time should be allocated for data cleanup and quality control.

**Reliability:** Paul, S. M. *et al*. ^1^estimate that lead optimization typically exhibits an 85% success rate. We find that the CRADLE-1 system exhibits a 90-95% success rate. We hypothesize that failure cases correspond to underlying biochemical limitations, as we have never observed failures in which a competing lead optimization attempt succeeded.

**Broad Generalizability:** the examples in Section 2 are selected to cover a wide diversity of protein optimization tasks. Spanning over a dozen modalities and over 50 distinct properties, we found the system capable of optimizing every target protein evaluated to date. This versatility suggests significant potential for novel applications - for example, black-box optimization of transcription factors or regulatory proteins using transcriptomic readouts as a fitness signal.

**Black box consumption of data:** sequence-function measurements are consumed without additional information such as the sequence or structure of a binding target. As an example, we speculate that this makes it possible to optimize a multi-step enzymatic pathway using only the readout of the final step.

**Low experimental throughput:** a single 96-well plate is sufficient data for each round. We do not routinely consider dataset sizes below 85 for multi-shot learning, as it becomes infeasible to run cross-validation to ascertain the quality of the models trained in Section 3. It is moderately common that mishaps occur when gathering wet lab data, and such validations are important for the robust function of CRADLE-1. Data coming from a single 96-well plate has been reliably observed to produce good results. This is consistent with the broader machine learning literature, in which sufficiently powerful foundation models may be refined in-domain with few samples (‘few shot learning’)^35–37^.

**Escaping the sunk cost fallacy:** teams often persist with stubborn campaigns past the point of diminishing returns due to the lack of an actionable stop signal. CRADLE-1 addresses this by providing a quantitative estimate of “optimization headroom”, computed from the ratio of sequences predicted to outperform the template and the magnitude of those predicted gains. In scenarios where this predicted improvement falls significantly short of the target profile - particularly in cases with sufficient data to train accurate models - it serves as a strong indicator that the campaign is unlikely to succeed. This allows teams to avoid unnecessary experiments on proteins that have effectively reached a biological ceiling.

### 4.1 lmplications

We consider a few possible key implications for organizational adoption of automated systems like CRADLE-1.

**Reduced operational expenditure:** from high reliability and multiple-factor improvements in speed.

**Adjustment of risk profiles:** as lead optimization improves in reliability, organizations will tolerate increased risk in other areas. For example, this makes it easier to pursue hard targets or rare diseases.

**Improved efficiency of capital allocation:** it is common for any individual drug development program to be paused and then resumed due to temporary budget constraints, etc., which imposes context-switching inefficiencies as work is resumed. This occurs because budget timelines are typically shorter than drug development timelines. However, faster lead optimization timelines may shorten drug development timelines to operate within budget timelines - for example, if novel pre-clinical candidates can consistently be obtained within a year.

### 4.2 Limitations

Wet lab assays must be good proxies for the properties requiring optimization. Properties like thermostability are straightforward to measure via techniques like differential scanning fluorimetry (DSF); however, properties like immunogenicity may require more sophisticated or costly assays such as MHC-associated peptide proteomics (MAPPs) or T-cell activation assays. CRADLE-1 does not provide a means of optimizing that which cannot be measured.

### 4.3 Extensions of CRADLE-1’s lead optimization

**Bispecific antibodies:** CRADLE-1 has been used to optimize bispecific antibodies, by performing two parallel runs of the system, corresponding to the variable regions of each arm whilst scFv-formatted. These are then reformatted and heterodimerized with knob-in-hole CH3 designs.

**Screentuning:** early-pipeline screening data may provide sequences without quantitative functional values. In such cases, we have successfully run CRADLE-1 using screening sequences in place of the multiple sequence alignment (Section 3); this is then used for evotuning and masking. Screening provides an alternative means for obtaining related sequences, especially in the CDRs, where the assumption of evolutionary conservation does not hold. On the other hand, if the enrichment values resulting from screening data are reliable, we successfully used the fold enrichments to fine-tune models predicting binding affinity (Appendix B.2.2).

**Identify template sequences:** early-pipeline screening data may provide large numbers of potential template sequences. In the same way as is used in CRADLE-1, diversity-aware ranking may be used to downselect these to a small and diverse set of template sequences.

## 5 Conclusion

We have introduced CRADLE-1, an automated machine learning system performing lead optimization of proteins. The system is able to reliably (>90%) achieve target product profile across all modalities tested (VHHs, scFvs, IgGs, peptides, enzymes, CRISPR systems, vaccines) and all properties (binding, activity, thermostability, aggregation, nonspecificity, immunogenicity,.), and do so faster (4–7×) than currently-typical alternatives.

### Commercial statement

The technical methods described in this paper are drawn primarily from Cradle’s machine learning workflows as of approximately two years prior to publication date. Publication at this time represents a trade-off between protecting commercial interests whilst upholding our responsibilities towards open science. For the avoidance of doubt, all results in this paper were still obtained using the methods described in this paper.

For anything to do with lead optimization of proteins, get in touch.

## Data availability

All data generated internally by Cradle, for the results discussed here, will be open-sourced in a future publication.

The results of Section 2.1 are already available on Proteinbase^38^.

## Supplementary Material

### A Experiments

#### A.1 Baselines

##### A.1.1 Anti-SARS-CoV-2 VHH Baselines

We evaluated the performance of CRADLE-1 against two distinct benchmarks. The first is a conventional protein engineering baseline consisting of a deep mutational scan of the CDR3 region (residues 98–108), followed by combinatorial library design. The initial CDR3 scan of 209 variants yielded three “hits,” defined as sequences exhibiting improved *T*_*m*_, expression, and affinity (*K*_*D*_) relative to the wild-type template. To progress these variants, we combined the three primary hits with the top five performers for each individual property to generate a second-round library of double mutants. This recombinatorial process was repeated for Round 3.

The second baseline is a computational pipeline utilizing ProteusAI^28^ implemented on top of an ESM-2-650M model. In the zero-shot phase (Round 1), we used the ESM-2-based generator to produce 192 sequences with the highest log-likelihood ratios relative to the template. For subsequent supervised rounds, we employed the ProteusAI MLDE module, training regressors across multiple architectures (Random Forest, Ridge, Gaussian Process, and SVM). Random Forest (seed 145) was selected as the optimal regressor for sequence selection. Candidates were generated using an exploration-exploitation factor of 0.5, with *T*_*m*_ as the primary objective, as the platform does not natively support multi-property optimization.

The cradle-1 optimization campaign was initiated with a zero-shot diversification of the template. In Round 2, the primary objective was shifted to maximize thermal stability (*T*_*m*_), while enforcing non-inferiority constraints for expression yield and binding affinity relative to the template. Although this round successfully identified several variants with stability gains exceeding 10°*C* over the wild-type, these candidates exhibited only moderate affinity toward the Omicron variant (double-digit nM *K*_*D*_). To address this, Round 3 was designed to prioritize Omicron affinity while maintaining a minimum *T*_*m*_ threshold of 65°*C*. This multi-objective refinement proved highly effective, yielding several elite candidates that achieved single-digit nM affinity against both targets while retaining a *T*_*m*_ above 70°*C*. This trajectory highlights the system’s ability to perform “active” Pareto frontier exploration-strategically shifting the population toward the optimal quadrant through successive, model-guided iterations.

###### A.1.1.1 Comparative Analysis

Results are presented in Figure 7. To facilitate comparison across multi-dimensional objectives, we visualize bi-objective Pareto frontiers, highlighting non-dominated variants within a 2D subspace. To ensure functional relevance, variants with expression levels below the template threshold were excluded from the analysis.

The results in Figure 7 (right) indicate that cradle-1 efficiently explores the functional landscape as early as the first round, effectively managing the trade-off between *T*_*m*_ and affinity. By Round 2, fine-tuned models successfully shift the population diagonally toward the upper-right quadrant. By Round 3, the integration of fine-tuned generators and models enables a significant leap in landscape modeling, resulting in a dense clustering of variants in the high-performance quadrant.

In contrast, the ESM-2-based ProteusAI pipeline (Figure 7, middle) identifies promising candidates in Round 1 but fails to maintain this trajectory in later rounds. We hypothesize that the lack of generator fine-tuning prevents the model from extrapolating beyond the natural sequence distribution. Furthermore, 76% of the generated sequences were lost due to poor expression. While the CDR3 scanning approach (Figure 7, left) preserves function effectively in the initial round, the combinatorial stacking of mutations produces only marginal improvements, highlighting the limitations of classical single-property greedy optimization compared to the ML-guided multi-property search space exploration of cradle-1. Table 1 summarizes the quantitative metrics for the top three variants from each methodology. Here, top candidates are defined as the most thermally stable variants that maintain binding affinities equivalent to or better than the wild-type template. Notably, the leading cradle-1 variant achieves a melting temperature more than 8°G higher than the next-best sequence identified by either baseline method.

**Table 1:** Top-ranking VHH candidates for multi-objective optimization. Variants are ordered by melting temperature (*T*_*m*_). Only the top three sequences per method that maintain or improve upon the template’s binding affinity are shown.

| Method | Tm | Kd (WT) | Kd (Omicron) | Expression |
| --- | --- | --- | --- | --- |
| Cradle | **71.4°C** | 5.28e-10 | **6.08e-09** | **0.17** |
| Cradle | 70.9°C | **1.86e-10** | 1.14e-08 | **0.17** |
| Cradle | 70.8°C | 5.04e-10 | 1.25e-08 | **0.17** |
| OSS ESM Method | 63.3°C | 1.52e-09 | 3.32e-08 | 0.05 |
| OSS ESM Method | 61.5°C | 1.54e-09 | 2.86e-08 | 0.15 |
| OSS ESM Method | 61.2°C | 8.74e-10 | 2.15e-08 | 0.09 |
| CDR DMS + Stacking | 60.9°C | 1.95e-09 | 9.79e-09 | 0.10 |
| CDR DMS + Stacking | 58.3°C | 1.94e-09 | 1.06e-08 | 0.09 |
| CDR DMS + Stacking | 58.1°C | 1.74e-09 | 1.40e-08 | 0.07 |

##### A.1.2 Anti-Her2 scFv Baselines

We further assessed the performance of the cradle-1 system through a comparative study on an anti-Her2 scFv. We used an scFv version of Trastuzumab as initial template. Given the high baseline affinity of the template (5nM), the primary optimization objective was set to improving *T*_*m*_ while maintaining expression and binding affinity above the template. The conventional protein engineering baseline comprised a comprehensive alanine scan across all six complementarity-determining regions (CDRs) to identify residues permissive to mutation. Since the initial scan yielded no direct “hits” (defined as variants with superior *T*_*m*_, expression, and affinity relative to the template), we established permissive criteria based on maintaining at least 60% of template expression, a *K*_*D*_ within 10-fold of the template, and a *T*_*m*_ within 1°*C* of the template. Six residues were identified as permissive. We subsequently performed site-saturation mutagenesis at these loci—excluding cysteine (to avoid non-native disulfide formation), alanine, and the template residue, generating a second-round library of 102 single-point mutants (17 * 6). In the third round, the top-performing mutations were combinatorially assembled on the lead sequence.

The computational baseline utilized the ProteusAI pipeline^28^ with an ESM-2-650M backbone. The zero-shot configuration (Round 1) mirrored the SARS CoV2 VHH study. For the supervised phase (Round 2), the MLDE module was trained on *T*_*m*_ using 59 expression-validated sequences. A Random Forest architecture (seed 145) was selected as the optimal regressor, generating 147 double-mutant candidates with an exploration-exploitation factor of 0.5. Round 3 followed an identical supervised learning framework with the additional data from Round 2.

###### A.1.2.1 Comparative Analysis

The results for all three methodologies are detailed in Figure 9. The initial cradle-1 diversification of the template sequence (Round 1) resulted in only 39 expression-validated sequences. To mitigate this “cold-start” challenge, we prioritized expression as the primary objective in Round 2 while applying an in silico constraint to ensure predicted *T*_*m*_ values exceeded the template. In Round 3 the primary objective was set to improving *T*_*m*_.

**Figure 9:**
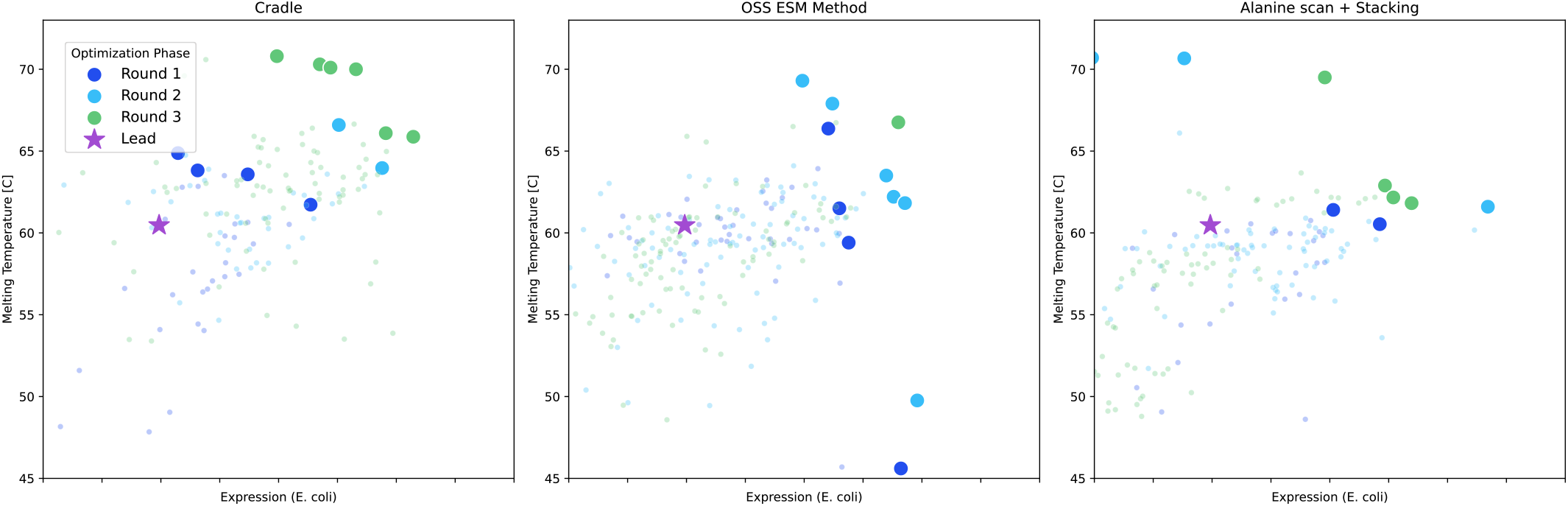
Comparative Pareto frontier analysis for multi-objective scFv optimization. Successive optimization rounds for an anti-Her2 scFv are visualized across three methodologies: cradle-1 (left), an open-source ESM-2-based method (middle), and alanine scan+CDR-DMS with stacking (right). Plots illustrate the trade-off between expression yield (*E. coli*) and thermal stability (*T*_*m*_). Successive cradle-1 rounds demonstrate a clear shift toward the high-performance upper-right quadrant.

Despite the sparsity of the Round 1 training data, the Round 2 cohort exhibited a significant improvement in expression yield (Figure 9, left, cyan markers). This enhanced baseline enabled a pivot back to *T*_*m*_ as the primary optimization objective for the final round. With only a single round dedicated specifically to thermal stability, cradle-1 successfully identified several variants with *T*_*m*_ values above 70°*C*, representing a 10°*C* improvement over the template. In comparison, both the classical and ESM-2-based baselines were unable to efficiently navigate the trade-off between expression and stability (Figure 9, middle/right). While the performance gap between cradle-1 and the baselines was narrower than observed in the SARS-CoV2 VHH campaign (reflecting the increased maturity of the Trastuzumab template), cradle-1 was the only system to demonstrate a controllable, directional expansion of the Pareto frontier toward the optimal quadrant across successive rounds. In contrast, the baseline distributions remained largely irregular and stochastic, suggesting their few high-performing variants were the result of chance sampling rather than an effective exploration of the functional landscape.

#### A.2 Ablations

##### A.2.1 MSA search database

We compared the following target databases:

1. **ColabFold:** Search with UniRef30, expand with BFD^39^ and Mgnify^40^.
2. **UniRef100:** Combines identical sequences and sub-fragments with 11 or more residues from any organism into a single UniRef entry (from UniProt).
3. **UniRef90:** Clusters UniRef100 sequences such that each cluster has sequences with at least 90% sequence identity to, and 80% overlap with, the the seed sequence.
4. **UniRef50:** Clusters UniRef90 seed sequences such that each cluster has sequences with at least 50% sequence identity to, and 80% overlap with, the seed sequence.

We evaluate the impact of database choice by evotuning models on MSAs built from searches against each database, and computing the Spearman rank correlations between model pseudo-log-likelihoods and P450 activity (Section 2.4) across a selection of rounds, see Figure 10. On this basis we selected the ColabFold database as our default choice.

**Figure 10:**
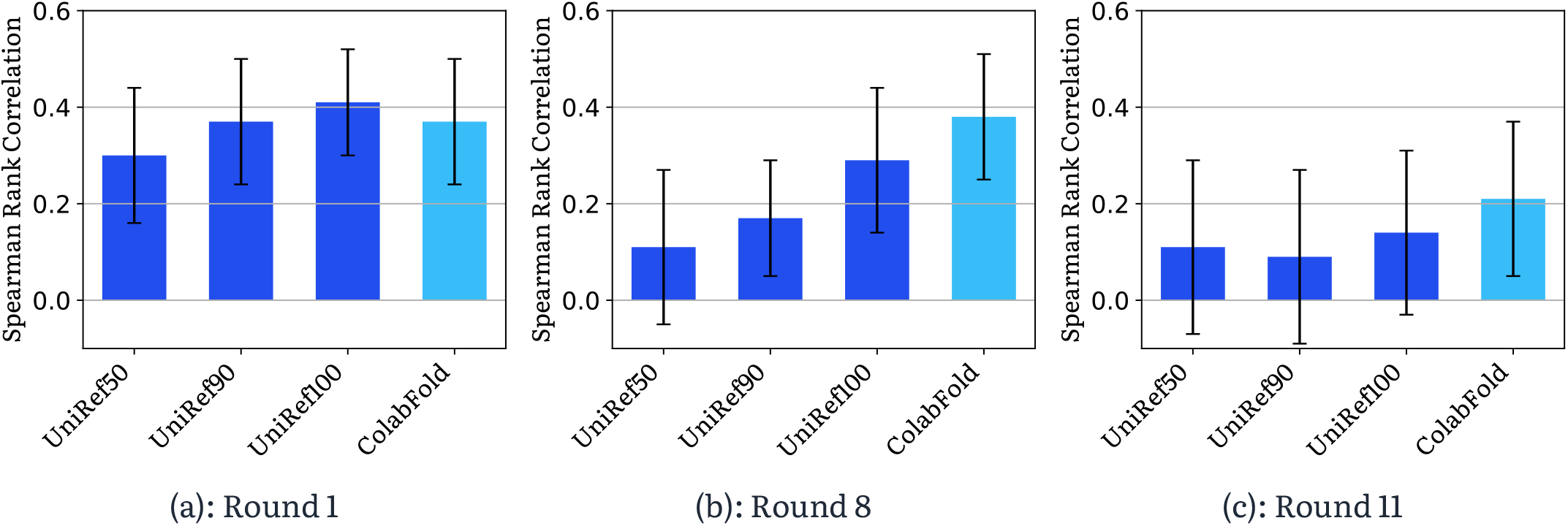
Spearman rank correlations between model pseudo-log-likelihoods and P450 activity, as choice of reference dataset varies.

##### A.2.2 MSA E-value filtering

The E-value is the expected number of matches of equal or better score that would be observed by chance given the database size. Lower E-values indicate more statistically significant matches. We ablated the E-value threshold used to filter MSA hits, varying how strict the search is.

We perform a sweep over the following E-values: 10^−300^, 10^−250^, 10^−200^, 10^−150^, 10^−100^, 10^−80^, 10^−65^, 10^−50^, 10^−25^, 10^−10^, 10^−1^]. See Figure 11.

**Figure 11:**
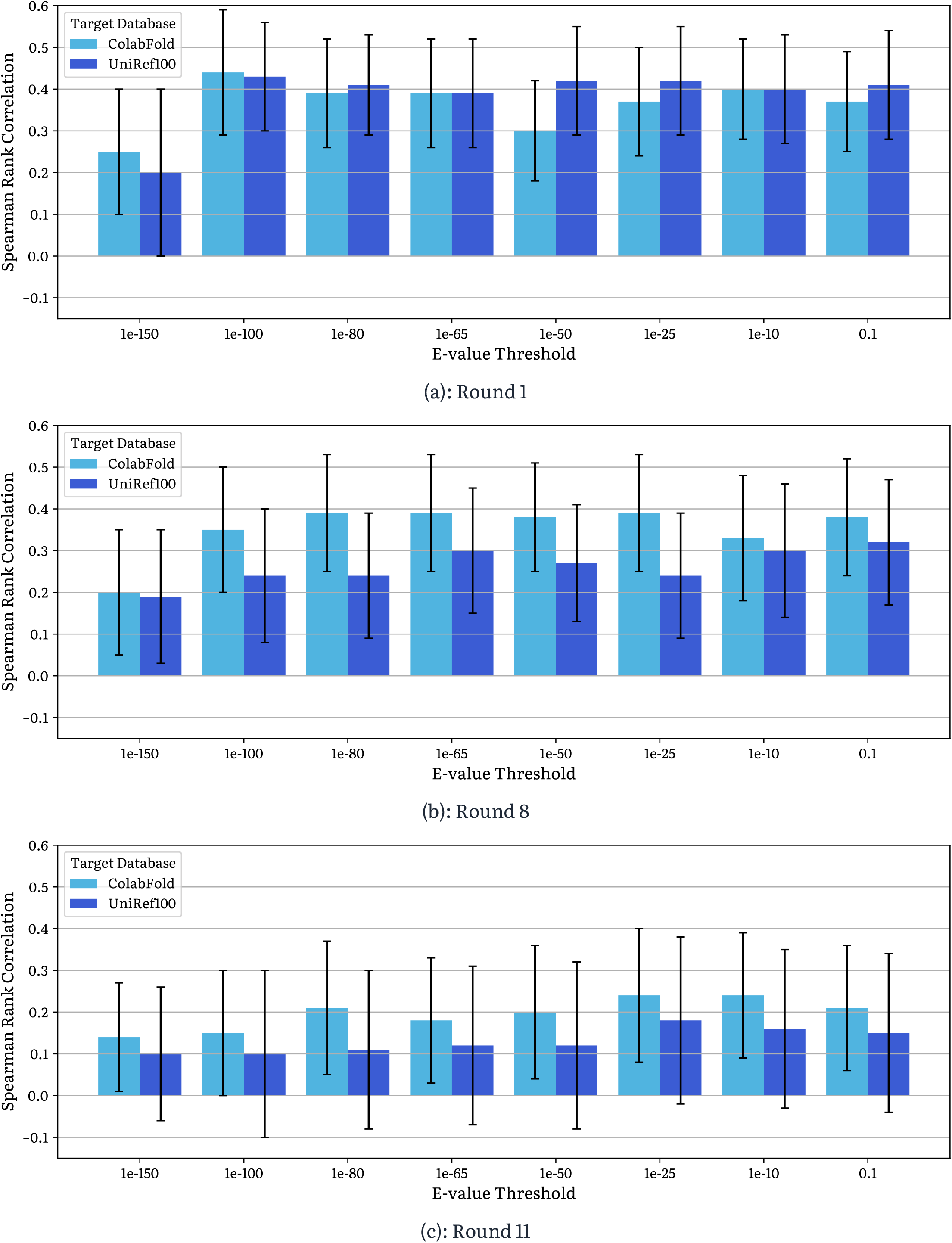
Spearman rank correlations between model pseudo-log-likelihoods and P450 activity, as E-value and choice of dataset varies. We consider both the ColabFold and the Uniref100 datasets. Whilst ColabFold consistently outperforms UniRef100, we do not observe any change based on E-value.

We find that no significant effect is observed with setting different E-value thresholds.

We find that the MSA dataset size has a measurable impact on performance (data not shown). To ensure our models are provided with sufficient evolutionary signal, we prioritize maximizing sequence retrieval. Consequently, we select a threshold of 10^{−1}^, which typically yields a sufficient number of sequences across diverse queries, as a safe choice.

##### A.2.3 Evotuning

To evaluate the gain provided by evotuning, we compute the Spearman rank correlations between model likelihoods and P450 activity, across both the evotuned and non-evotuned (foundation) model.

We observe that evotuning provides a substantial improvement. See Figure 12.

**Figure 12:**
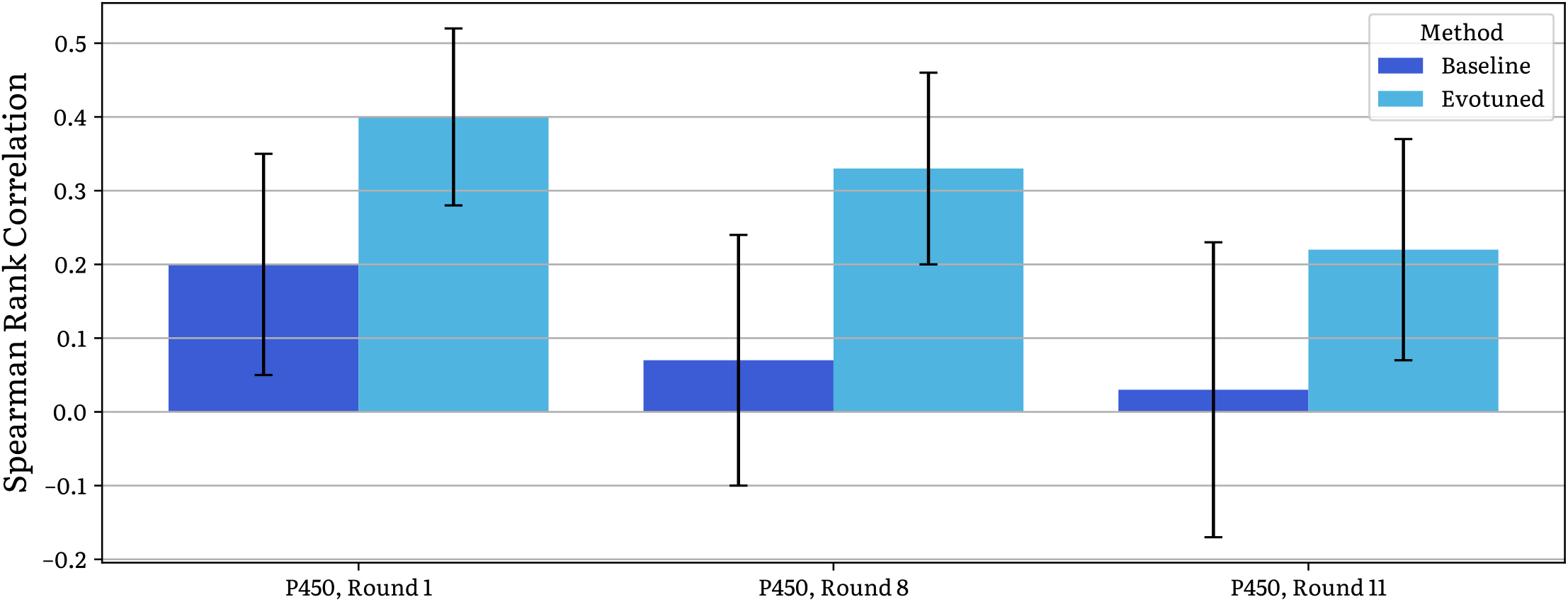
Spearman rank correlations between between pseudo-log-likelihoods and P450 activity, across three rounds. Evotuned models consistently achieve higher correlations than the foundational baseline.

##### A.2.4 g-DPO

To isolate the effect of DPO vs. evotuning, we evaluate the (i) Spearman rank correlation between model pseudo-log-likelihoods and experimental assay measurements, and (ii) the quality of generated sequences.

DPO consistently improves rank correlation over the evotuned reference model across datasets, indicating better alignment between model likelihoods and experimental measurements. g-DPO matches or slightly exceeds the performance of standard DPO while providing substantial training speedups. When evaluating generated sequences, both DPO and g-DPO produce designs with improved predicted properties relative to the evotuned reference, which confirms that preference-based optimization captures task-specific signal beyond evotuning alone. See Figure 13.

**Figure 13:**
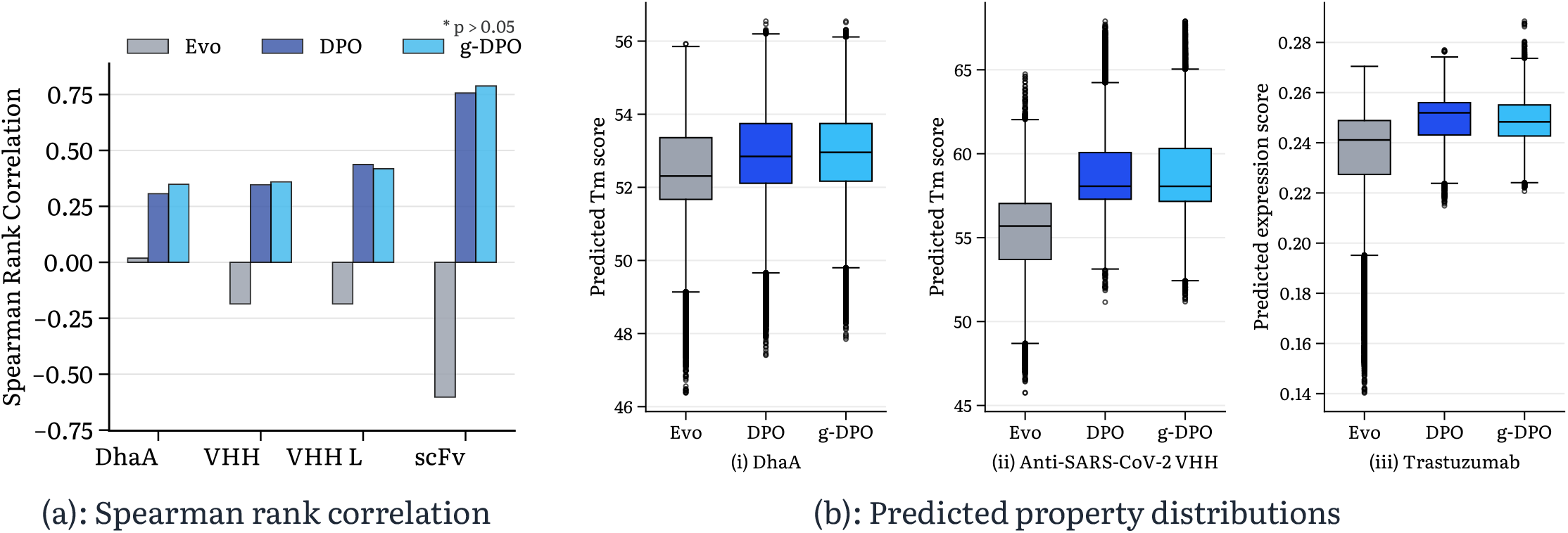
In silico evaluation of DPO. (a) Spearman rank correlation comparing performance of evotuned, DPO, and g-DPO models. (b) Predicted property distributions of sequences generated via beam search. Kolmogorov-Smirnov (KS) tests confirm that g-DPO and DPO yield nearly identical distributions, both showing significant improvement over the base reference model.

##### A.2.5 Hyperparameters

Substantial ablations of hyperparameters were performed through *in silico* evaluation of results. For brevity these ablations are not shown here.

The hyperparameter optimization procedure was performed manually and without reference to Bayesian optimization tools. The focus was on finding an end-to-end system that reliably works across all modalities considered; more careful hyperparameter tuning is a focus of future work.

### B Further results

#### B.1 scFv framework optimization for binding to a transmembrane protein

The original Adaptyv Bio EGFR competition allowed up to ten entrants per competitor; as a practical matter only one cradle-1-derived sequence was selected, which was the one marked as ‘Cradle 4’ in Figure 2(a). The other sequences were later generated under the same conditions and without any additional model training.

Only a single round of wet lab validation is performed, and as such the only input data is evolutionarily-related data, without in-context sequence-function data.

As this project involved framework mutations, we additionally perform an *in silico* evaluation of immunogenicity risk.

We use the LakePharma Antibody analyzer^41^ to calculate a T20 score, which quantifies how closely an antibody’s variable region resembles human antibodies. The predicted scores indicate that our mutations did not reduce the humanness of the variants.

We use NetMHCII^42^ to predict the binding affinity of each peptide within the sequence to different HLA-DR, HLA-DQ, and HLA-DP alleles. The number of core peptides binding to MHC is determined by counting the unique core peptides that rank within the top 2% of 1,000,000 randomly chosen interactions. Predictions indicate that we did not introduce any mutations that could increase the risk of unwanted immunogenic responses.

See Figure 14.

**Figure 14:**
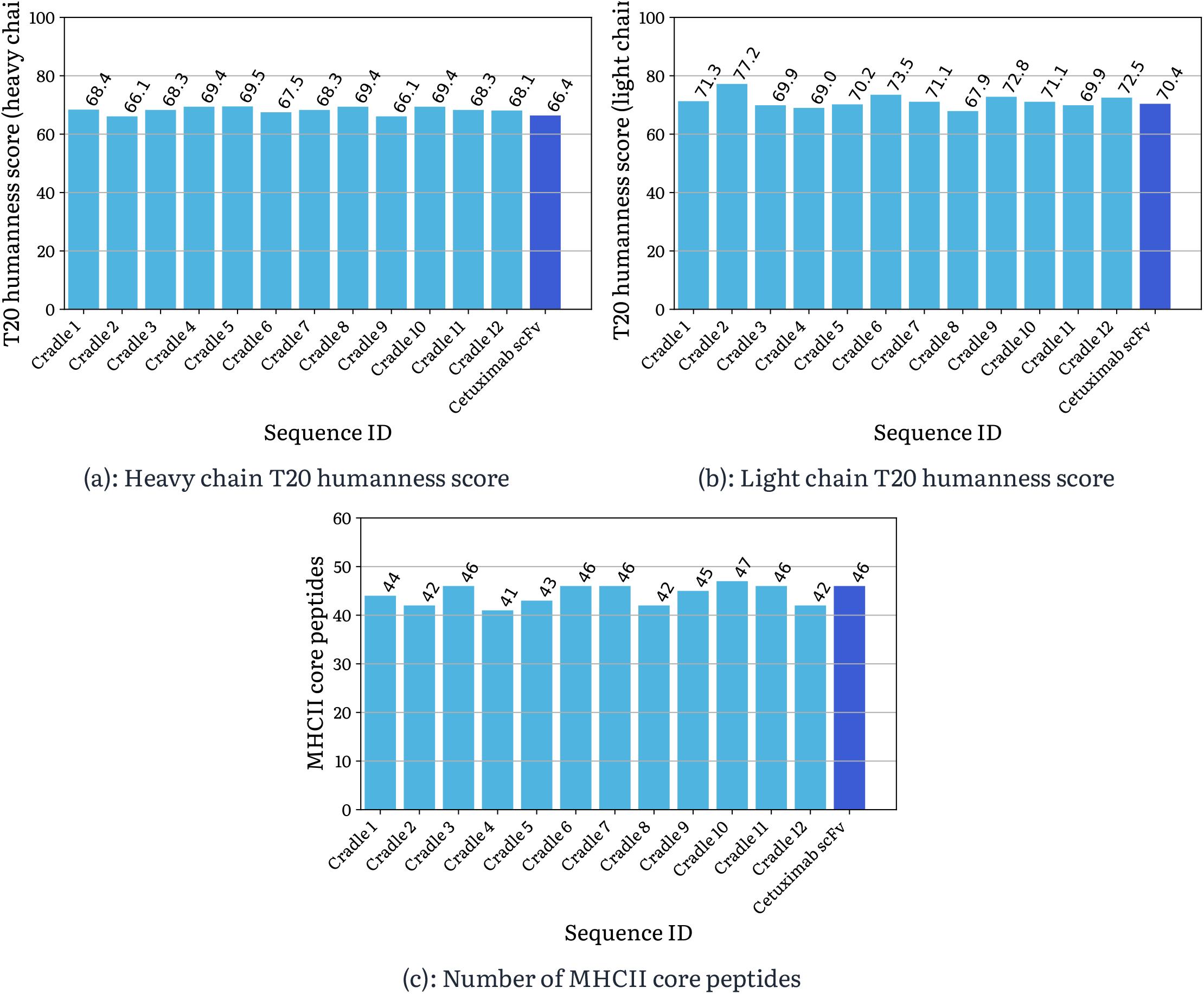
*in silico* predicted immunogenicity of cradle-1-generated sequences, and template Cetuximab. Top panels: humanness score for heavy and light chains; higher is better. Bottom panel: MHCII core peptides; lower is better. The template is shown in a different color. Little difference is seen between generated candidates and the template.

All sequences are available on Proteinbase.^38^

#### B.2 VHHs with picomolar and polyspecific binding

##### B.2.1 SARS-CoV-2

The template sequence is **MQVQLVETGGGLVQPGGSLRLSCAASGFTFSSVYMNWVRQAPGKGPEWVSRISPNSGNI-**

**GYTDSVKGRFTISRDNAKNTLYLQMNNLKPEDTALYYCAIGLNLSSSSVRGQGTQVTVSS.** The chosen ‘best’ sequence is **MEVKLQESGGGLVQPGGSLRLSCAASGFTFSSVYMNWVRQAPGKGPEWVSRISPDGGSTYYADSVKGRFTISRDNAKN-**

**LLYLQMNNLKPEDTALYYCAIGLNLSSSSVRGQGTQVTVSS**, and this is 11 mutations away from the template sequence.

The first round was run in ‘zero shot’/’diversify’ form without sequence-function data. The second round was run with melting temperature as the primary objective. The third round was run with binding to Omicron as the primary objective.

We plot the data for all pairs amongst the four properties, across all three rounds of optimization, in Figure 15. Each round of wet lab iteration progressively pushes out the Pareto frontier for all pairs of properties.

**Figure 15:**
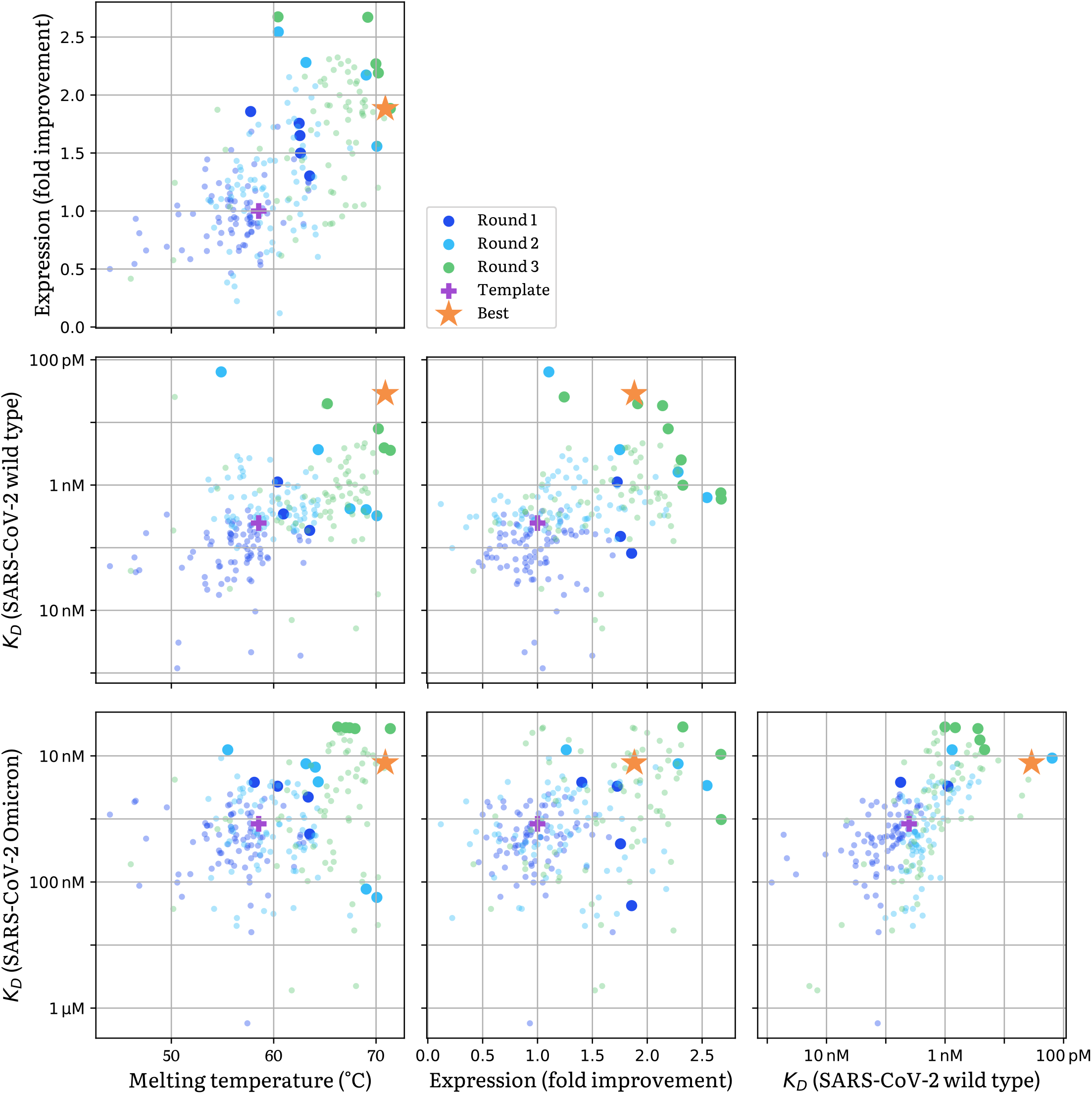
VHHs obtained with cradle-1 for polyspecific binding to SARS-CoV-2 wild type and SARS-CoV-2 Omicron. All pairs of properties are shown. There are four properties in total (leading to six pairs, corresponding to the six panels of the figure), and data is drawn from three rounds of optimization. Being higher or to the right is better. The two-property Pareto-optimal candidates in each round are highlighted. The chosen ‘best’ candidate is shown as an orange star. The original template sequence is shown as a pink plus.

Picomolar binding values are reported despite the lowest SPR analyte concentration being 977 pM. The true precision of this number was the topic of some debate amongst our biology team; eventually it was decided that the reported numbers were still reasonable.

##### B.2.2 Snake venom

Two rounds of optimization are performed. The first is run as a diversify/zero-shot round, and the second is run with thermostability as the primary objective, whilst setting binding and expression as relative constraints (tasked to improve relative to the template sequence).

Figure 16 shows fold-improvement in binding, and the expression against thermostability for cradle-1-generated VHHs.

**Figure 16:**
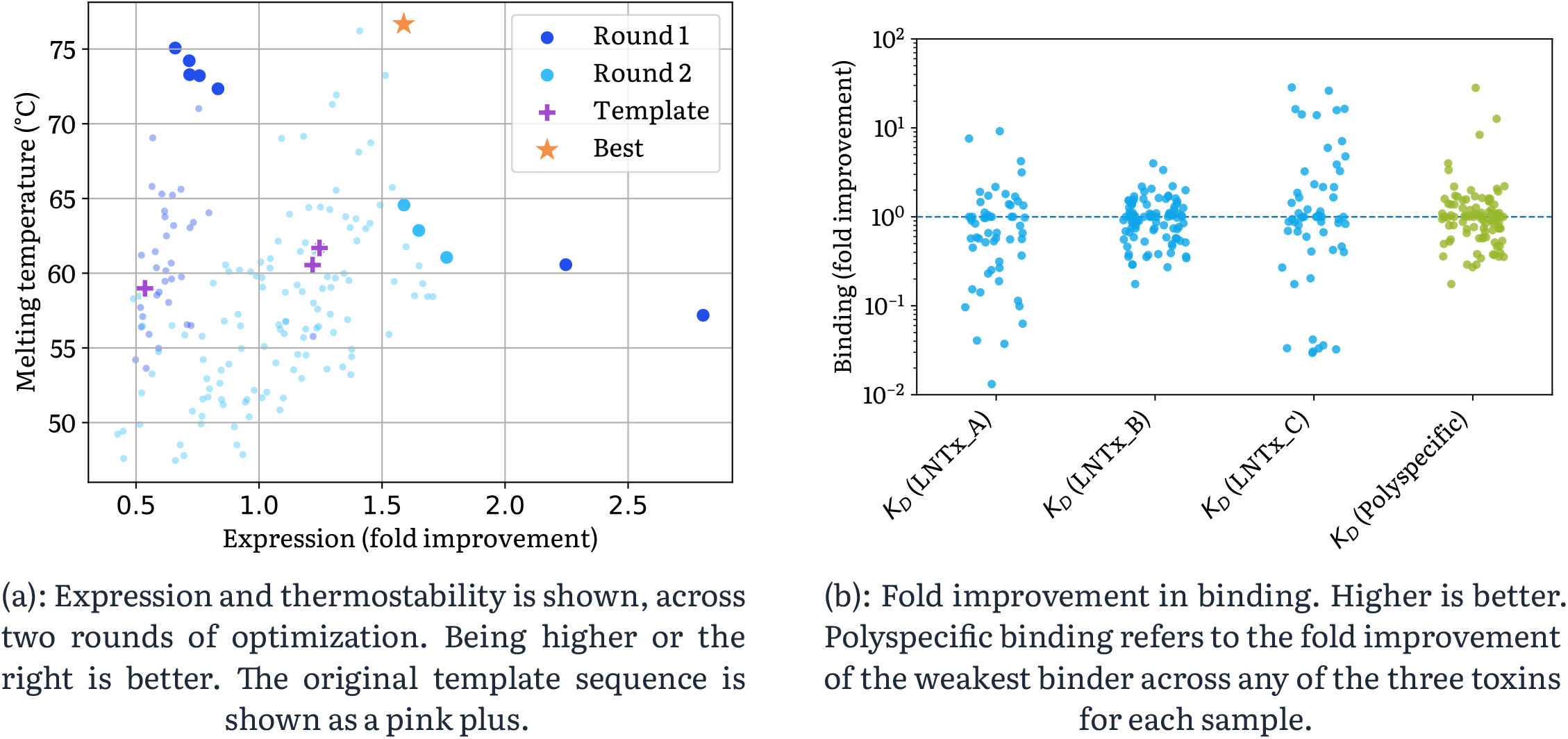
VHHs obtained with cradle-1 for polyspecific binding to three snake venom toxins. Polyspecific binding is calculated as 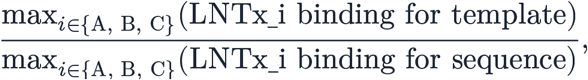, reflecting the polyspecific improvement rather than the worst single improvement.

#### B.3 Multi-property optimization of a bispecific VHH

We used the optimization of a bispecific VHH to evaluate the ability of the cradle-1 system to learn from partial data in sequence-space: single-domain, unlabeled screening data, and fully characterized variants for the individual domains were provided to the cradle-1 pipeline as an ablation study to generate three separate libraries for in-vitro testing. While the no-context approach, providing no single-domain data, was able to generate a function-preserving set of designs with improved binding for each individual target, the context-aware approach was able to generate improved variants more reliably, as demonstrated by the distribution shift shown in Figure 17. Additionally, the use of labeled data resulted in variants with a larger nominal improvement over the templates with a controllable trade-off over properties as indicated by the defined constraints. In this experiment, screening data were reduced to sequence information: no read counts or enrichment values were provided to the model. The lower information content of these data is reflected in the limited improvement relative to the labeled data approach. Our goal was to simulate different starting points that are common when engaging in a bispecific binder optimization campaign. Often, some form of single-domain information is available. Extracting salient information from these data can have a significant impact on the quality of the generated library. We explicitly chose targets for which we had previously run both hit-identification and lead-optimization campaigns to demonstrate the utility of the different data types. The templates were chosen from the single-domain campaigns, and an initial hit-identification over 16 candidates was performed to select the most promising configuration (the order of the VHH in the full chain and the linker length) before proceeding to the optimization rounds documented here.

**Figure 17:**
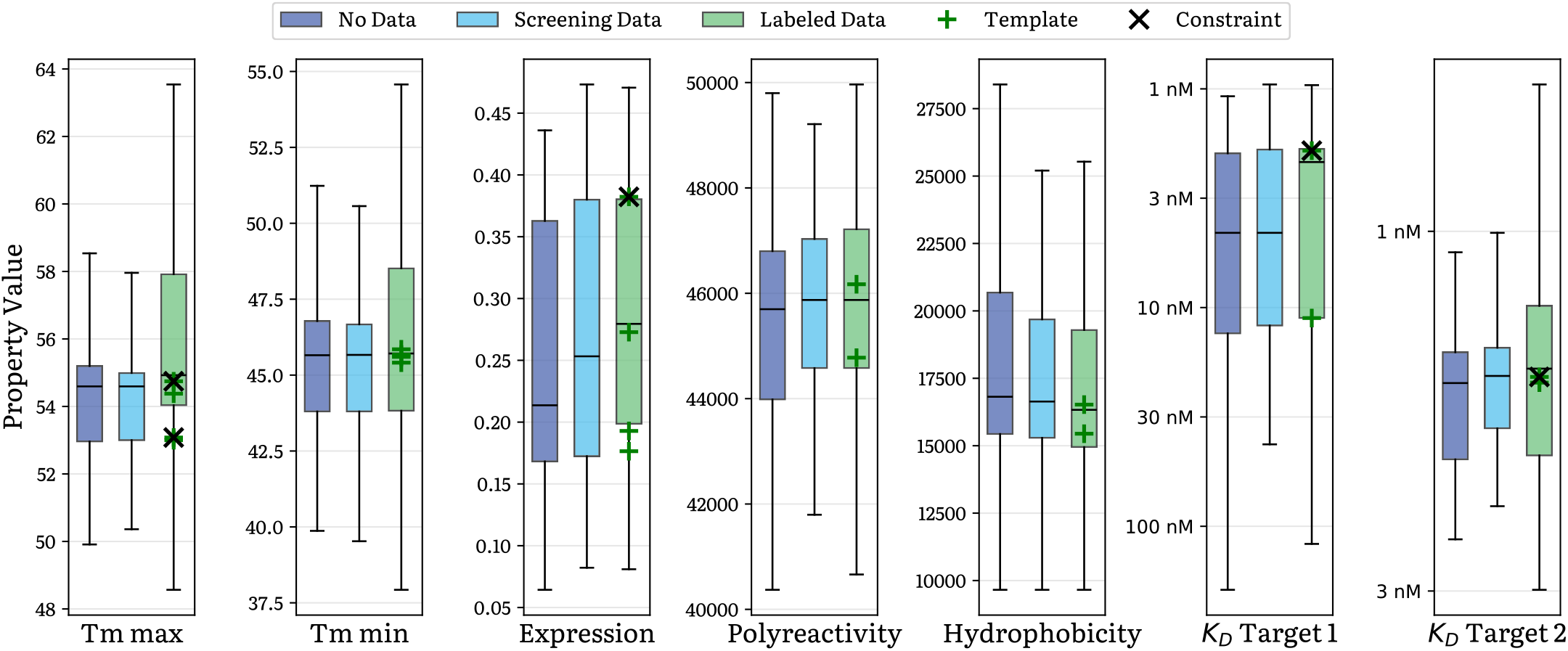
Distribution of assay values for all characterized properties across different design strategies of a bispecific VHH. While cradle-1 can generate function-preserving variants using only the template sequences as a starting point, the quality of the generated library is significantly improved when adding partial context data, such as screening data for the individual domains, or labeled plate-based data from previous single-domain engineering campaigns. Labeled data were available for all properties except hydrophobicity and polyreactivity.

#### B.4 Joint optimization of thermostability and expression of a haloalkane dehalogenase

The first round of optimization was a diversify/zero-shot round. Observing that some variants had poor expression, the second round was run with thermostability as the primary objective whilst setting expression as a relative constraint (to improve relative to the template sequence).

The final best sequence was manually selected as the one with the best thermostability, as it also had excellent expression.

The starting sequence was **MSEIGTGFPFDPHYVEVLGERMHYVDVGPRDGTPVLFLHGNPTSSYLWRNIIPH-**

**VAPSHRCIAPDLIGMGKSDKPDLDYFFDDHVRYLDAFIEALGLEEVVLVIHDWGSALGFHWAKRNPERVKGIACMEFIR-**

**PIPTWDEWPEFARETFQAFRTADVGRELIIDQNAFIEGALPKCVVRPLTEVEMDHYREPFLKPVDREPLWRFPNELPI-**

**AGEPANIVALVEAYMNWLHQSPVPKLLFWGTPGVLIPPAEAARLAESLPNCKTVDIGPGLHYLQEDNPDLIGSEIARWL-PAL**, and the selected best sequence is **MSEIGTGFPFDPHYVEVLGERMHYVDVGPRDGTPVLFLHGNPTSSYLWRNI-**

**IPHVAPSHRCIAPDLIGMGKSDKPDLDYFFDDHVRYLDAFIEALGLEEVVLVIHDWGSALGFHWAKRHPERVKGIACME-**

**FIRPIPTWDEWPEFARETFQAFRTPDVGRELIIDQNAFIEGALPKCVVRPLTEVEMDHYREPFLKPADREPLWRFP-**

**NELPIAGEPANIVALVEAYMNWLHQSPVPKLLFWGTPGVLIPPAEAARFAESLPNCKTVPIGPGLHYLQEDNPDLIG-**

**SEIARWLAAL**, which differs by 6 mutations.

#### B.5 Optimizing activity of a P450 enzyme All round were run with thermostability as the primary objective, using available historical sequence-function data

Top-1 round improvement plots are calculated by computing the difference between the single best performers in each round, and then computing their violin plot. Top-5 round improvement plots compute the difference between the single best performers in each round, and the differences between the second-place performers in each round, *mutatis mutandis* the third-place/fourth-place/fifth-place, and then computing a violin plot of all such differences. Dashed lines indicate the median value.

#### B.6 Failure case: optimizing activity of a serine–pyruvate aminotransferase

We document a failure case of the cradle-1 system, for optimizing an alanine-glyoxylate aminotransferase enzyme (accession P21549), which also has activity as a serine-pyruvate aminotransferase. Three rounds of two-property optimization were performed, with the goal to optimize both thermostability and this secondary activity. Whilst the melting temperature was successfully optimized from 47.8 °C to 67.3 °C, the activity was optimized by only 1.96x. This activity improvement was considered insufficient to merit as a success.

To investigate whether this failure could have been avoided with some alternate approach, we challenged two experienced protein engineers (one identifying as a biologist, another identifying as a bioinformatician) to attempt the same problem; they were likewise each given three rounds of optimization. The best competing melting temperature was 73.3 °C and the best competing fold improvement of activity was 1.69x. This was likewise regarded as insufficient to merit a success.

Overall the Pareto frontier of candidates generated by the cradle-1 system seemed to outperform those provided by both protein engineers, even though the overall project did not succeed. See Figure 18.

**Figure 18:**
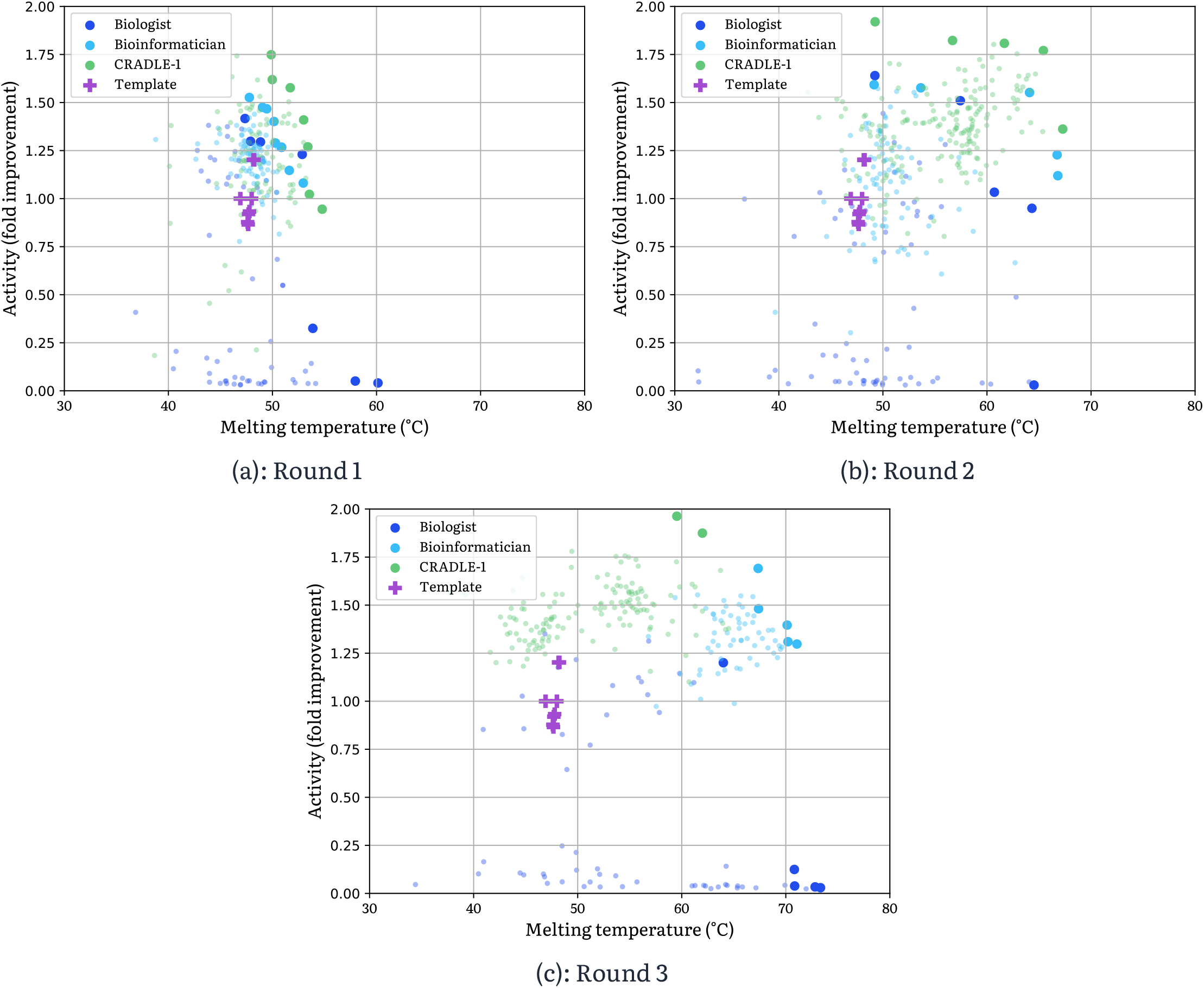
Activity and thermostability for a serine-pyruvate aminotransferase, across three rounds of optimization, by both cradle-1 and two protein engineers (‘biologist’, ‘bioinformatician’). Being higher or to the right is better. Pareto-optimal sequences (defined as those that are Pareto-optimal across that round and all previous rounds) are highlighted; the assayed properties of all other sequences are shown faded. The original template sequence is shown as a pink plus; it is shown multiple times corresponding to biological and technical replicates. Whilst the Pareto frontier of cradle-1-generated sequences pushes out round-over-round, and generally outperform those of the protein engineers, the maximal activity improvement is low and for this reason the project is considered a failure.

### C Wet lab protocols

We detail the wet lab methods for the results that were run internally at Cradle.

#### C.1 DNA design and synthesis

To prepare the designed protein variants for synthesis, a computational reverse translation workflow was employed. The process began with the amino acid sequences of the designed variants and their respective parental sequences. If specified, a hexahistidine tag was computationally added to either the N- or C-terminus of all sequences.

To handle insertions and deletions while preserving codon usage, all variant and parental protein sequences were first aligned using MAFFT43 (v7.526) to generate a multiple sequence alignment. A codon map was then constructed based on this alignment and the known DNA open reading frames (ORFs) of the parental sequences. For each position in the alignment, the codon used in the parental ORF was mapped to its corresponding amino acid. This mapping was then used to reverse-translate the aligned variant sequences.

For novel mutations or insertions not present in any parent, codons were selected from a standard *Escherichia coli* codon usage table. When a novel amino acid change at a specific position was encountered for the first time, a codon was chosen randomly, weighted by the natural codon frequency in *E. coli*. This codon choice was then recorded. To ensure consistency across the experiment, any other designed sequence featuring the identical amino acid change at the same position was assigned this same recorded codon. This method maintains the parental codon bias where possible, while ensuring that identical novel mutations are encoded by identical codons. The resulting DNA ORFs were then computationally flanked by user-defined 5’ and 3’ sequences to generate the DNA sequences ready to order at Twist Bioscience.

#### C.2 Cloning, transformation, and colony picking

Per round 384 gene fragments were ordered from Twist Bioscience, pooled together, and assembled into the expression vector pET11a via a pooled Gibson Assembly and transformed into *E. coli*.

The received gene fragments were first dissolved in 100 µL UltraPure Nuclease-Free water (Invitrogen, 10977035) to a final concentration of 100 ng/µL. After shaking for 30 minutes at 50°C and 1 000 pm to dissolve, 5 µL per well was aspirated and combined into a fragment pool. The Gibson assembly reaction was prepared using a ±10:1 molar ratio of fragments : linearized vector backbone pET11a-pT5-YFP (1 lacO) for a 10 µL reaction with NEBuilder^®^ HiFi DNA Assembly Master Mix (NEB, E2621). Any control or template sequences (hereafter called reference sequences) were assembled in a separate Gibson assembly reaction with the same components. Gibson assembly was run for 20 minutes at 50°C.

Per reaction mix, 5 µL was transformed into either SHuflle^®^ Express Competent *E. coli* (NEB, C3028) (for anti-SARS VHH and anti-snake venom VHH) or BL21 Competent *E. coli* (NEB, C2530H) (for haloalkane dehalogenase and serine-pyruvate aminotransferase libraries), following NEB’s instructions. For reference sequences, 5 µL of each reaction were transformed into the respective competent cells. After recovery, 100 µL of the transformants were plated on omni trays with LB (Gibco, 10855001) + 1% glucose (ThermoScientific, 450740010) + 50 µg/ mL carbenicillin disodium salt (FisherScientific, BP2648250) (LB/glu/carb) + agar and spread with glass beads to get even colony distribution. For templates and controls, transformations were streaked to get single colonies on individual Petri plates. Plates were incubated 16 hours at 37°C. If the plates did not show enough colonies (>4 fold the number of designed sequences) in this direct-to-expression host transformation, we went via a high-efficiency cloning host.

In such case, 5 µL of the Gibson assembly product was transformed into the higher-efficiency cloning host NEB13 10-beta Competent *E. coli* (High Efficiency) (NEB, C3019), in case the direct transformation into expression host did not generate a sufficient number of colonies. In this approach, the NEB^®^ 10-beta transformations were spun down after recovery, resuspended in approximately 50 µL of media, plated on Petri plates with LB/ glu/carb + agar, and spread with glass beads. After incubation at 37°C for 16 hours, the lawn of NEB^®^ 10-beta transformants were scraped up and resuspended in 1 mL of nuclease-free water and spun down into a pellet. Plasmid DNA from the cell pellet was extracted using the QIAprep Spin Miniprep Kit (Qiagen, 27104) following the manufacturer’s protocol. Plasmid DNA was quantified using the Qubit^™^ dsDNA BR Assay Kit (Invitrogen, Q32850). 30 ng of plasmid DNA was then transformed into the respective expression host (SHuflle^®^ Express Competent *E. coli* or BL21 Competent *E. coli* following the manufacturer’s protocol, again plating 100 µL of transformants on multiple omni trays with LB/glu/carb + agar.

Single colonies were picked from omni trays into 384-well deep plates containing 100 µL LB/glu/carb media using the PIXL automated colony-picker (Singer Instruments). The number of colonies picked was equal to 4x the number of designs (i.e. 1,536 colonies for a library of 384 designs). Individual colonies for templates and controls were hand-picked in duplicates into the final few wells of each plate. Plates were incubated at 37°C overnight, shaking at 1 000 rpm. The following day, 20 µL from each plate were stamped into sterile 384-well plates with 20 µL of 50% glycerol, and stored at -80°C.

#### C.3 Barcode PCR and sequencing

Since gene fragments undergo a pooled Gibson assembly and transformation, *E. coli* strains grown from picked single colonies must be sequenced to identify which sequences were built and cloned properly prior to expression. To match sequence reads to the original location of the corresponding strain in each 384-well plate, unique barcode pairs are assigned to each well and incorporated via barcode PCR.

Barcoding PCR reactions were prepared containing 2X repliQa HiFi ToughMix^®^ (Quantabio, 95200), 0.5 µL of the overnight culture, and 100µM for a total reaction volume of 10 µL. Plates were sealed and vortexed for 10 seconds and spun down before running PCR reaction on a thermocycler. The PCR reaction consisted of a pre-denaturation at 98.0°C for 2 minutes, followed by 32 cycles of (a) denaturation at 98.0°C for 10 seconds, (b) annealing at 58°C for 5 seconds, and (c) 68°C for 5 seconds/kb, with a final extension at 68°C for 2 minutes.

Barcoding PCR reactions were pooled (5 µL per well) and mixed. 40 µL of the PCR pool were cleaned up using the DNA Clean & Concentrator-5 kit (Zymo Research, D4004), and the final concentration was determined using the Qubit™ dsDNA BR Assay Kit as above.

The Ligation Sequencing DNA V14 Kit (Oxford Nanopore Technologies (ONT), SQK-LSK114) with the NEBNext^®^ Companion Module (NEB, E7180) was used for library sequencing preparation starting with 200 fmol of DNA from the barcode PCR pool, following the ONT protocol. Libraries were loaded onto MinION Flow Cells (ONT, R10.4.1) in MinION Mk1B devices per the manufacturer protocol and run using the MinKNOW application for 3 hours without base-calling.

#### C.4 Sequencing pipeline and hit selection

All sequencing data was processed using a custom bioinformatics pipeline developed in Python and orchestrated as a reproducible workflow. The entire software environment was containerized using Docker, based on a Debian 12.11 image.

Once at least 750 000–1 000 000 reads were obtained, the analysis workflow was initiated. Raw pod5 files were basecalled using Dorado (v0.5.2) with the `r1041_e82_400bps_sup_v420` super accuracy model. Reads shorter than 30 bp were filtered out using seqtk (v1.4 or above). Barcode sequences were identified using lastal (v1548 or above) and reads were demultiplexed into individual wells.

For each well, reads were aligned against all possible reference and input sequences (parental, negative control, and designed) using minimap2 (v2.26 or above) with the -x map-ont preset. A primary reference for each well was determined by identifying the single reference sequence to which at least 90% of the reads mapped. Wells were flagged for failure if this criterion wasn’t met.

For wells that passed this initial mapping QC, variant calling was performed against all the well reads mapping to their assigned primary reference using Clair3 (v1.2.0) with the `-platform=11ont11` flag. The resulting variants underwent a final QC step. Wells were disqualified if any variant was detected with an allele frequency (AF) between 0.2 and 0.7, as this indicates a mixed population. Mutations with an AF > 0.7 were accepted and applied to the reference sequence to generate the final consensus sequence for the well. Variants with an AF < 0.2 were considered sequencing noise and ignored.

From these, designs (including all templates and controls) that passed QC were mapped to new destination wells in 96-well plate format. Using a Tecan Fluent, 0.5 µL of the “hits” were picked from the source wells of the 384-well glycerol-stocked plates into new 96-deep well plates with 900 µL of LB/glu/carb.

#### C.5 Growth and expression

Hit-picked culture plates were first incubated at 37°C for 16-20 hours while shaking at 1 000 rpm. 2.5 µL per overnight culture were stamped into new 96-deep well plates with 900 µL LB/glu/carb. Incubation continued at 37°C for 6 hours while shaking at 1 000 rpm. After 6 hours, 36 µL of seed culture were inoculated into fresh 96-deep well plates with 900 µL per well of pre-prepared expression media using the EnPresso^®^ B expression system (EnPresso, B11001) per the manufacturer’s protocol. Briefly, 4 tablets of EnPresso^®^ B (03EBX221, EnPresso) per 100 mL of sterile water were dissolved, before adding 50 µL of Reagent A (EnPresso) per 100 mL of water, plus carbenicillin up to 50 µg/mL, and aliquoting 900 µL of this media into each well. Incubation continued at 30°C for 18 hours at 1 000 rpm.

After 18 hours, induction media was prepared by dissolving 2 tablets of EnPresso13 Booster (03EBT221, EnPresso) per 10 mL of sterile water, supplemented with 50 µL per 10 mL media and a final concentration of 10 mM IPTG (Thermo Scientific^™^, R1171). 100 µL of induction media was added to each well, and incubation continued for 24 hours at 30°C for 1 000 rpm. After 24 hours, cultures were spun into pellets by centrifuging at 3 000 x g for 3 minutes, and plates were decanted to remove culture media.

#### C.6 Lysis

Lysis buffer was prepared with a final concentration of 1X FastBreak™ Cell Lysis Reagent (Promega, V8573), 30 U/mL DNAse I (Roche, 10104159001), supplemented with cOmplete™, EDTA-free Protease Inhibitor Cocktail (Roche, 11873580001) per manufacturer’s instructions. 803 µL of this solution were dispensed per well, followed by mixing rigorously by pipetting up and down to resuspend the pellet. Plates were shaken for 40 minutes at 450 rpm to lyse the cells. After 40 minutes, NaCl was added to a final concentration of 500 mM, mixed in by briefly pipetting up and down. Plates were centrifuged for 10 minutes at 3 000 x g at 4°C to pellet the debris. At this stage, a small sample of clarified (lysed cell debris centrifuged down to pellet) lysate (±30 µL) was saved in another plate for lysate-based assays.

#### C.7 Expression assay

To measure expression, 2 µL of each clarified lysate was analyzed with the LabChip^®^ GXII Touch™ protein characterization system (Revvity). The HT Protein Express LabChip (Revvity, 760499) and Protein Express Assay Reagent Kit (Revvity, CLS960008) were used to prepare and run samples, per the manufacturer’s protocol. Expression data was analyzed, briefly, by setting a 4 kDa window around the known size of the protein of interest and calculating the area-under-the-curve (AUC) for that region minus the AUC for the entire lysate, to give a relative measurement of expression over background.

#### C.8 Binding kinetics assays

For antibody fragments (anti-SARS VHH, anti-snake venom VHH), binding kinetics measurements were measured by SPR on the Carterra LSAXT using a SAHC30M chip. The chip was first conditioned with 1 minute injections of 10 mM glycine pH 2.0, 1X HBSTE (10 mM HEPES pH 7.4, 150 mM NaCl, 3 mM EDTA, 0.05% Tween-20) running buffer, and 20 mM NaOH. Next, an anti-his capture lawn was prepared by injecting a prepared solution of 75 ng/µL biotinylated THE™ His Tag Antibody (GenScript, A00613) over the sensor chip for 10 minutes. In general, a capture level of ±2 000 RU for the anti-his lawn was achieved.

For anti-SARS VHH proteins, SARS-CoV-2 spike recombinant binding domain (R&D Systems™, 10542CV100) and SARS-CoV-2 BA.2 spike recombinant binding domain (R&D Systems™ 11148CV100) were used as antigens. For anti-HER2 scFvs, ErbB2 recombinant protein (Invitrogen™, A42494) was used as an antigen. For anti-snake venom VHH, 3 different 3-finger toxins (named LNTx_A, LNTx_B, and LNTx_C) were used as antigens. Antigen dilutions were prepared in 1X HBSTE + 1% PEG3350. When antigens contained high refractive index (RI) stabilizers in the commercial product such as glycerol or trehalose, antigens were buffer exchanged using Vivaspin columns (Cytiva, 28932235 or 28932236) per the manufacturer’s instructions. Antigens after buffer exchange were quantified by bicinchoninic acid (BCA) assay (Merck, 71285-3) against a BSA standard curve. Antigen concentrations were optimized for each antigen (starting concentration from 250 nM-1 uM, serial dilution factor of 1:3 to 1:5), for a total of 6 antigen concentrations for each run.

Capture kinetics experiments were performed using a running buffer of 1X HBSTE + 1% PEG3350 with the chip at 25°C. Variant libraries, prepared in a dilution of anywhere from 1:20-1:200 in 1X HBSTE depending on the ideal capture level for kinetics (generally, 150-200 RU to achieve Rmax of ±50-100 RU) in deep 384-well plates, were captured on the chip for 5 minutes. 5 injections of 1X HBSTE + 1% PEG3350 were first applied to achieve a stable baseline, followed by injections of the antigens in ascending concentrations, using a 5 minute association and 10 minute dissociation. After all concentrations were injected, the chip surface was regenerated by 3 × 30 second pulses with 10 mM glycine pH 2.0.

Raw kinetic data were analyzed using the Carterra Kinetics software 1.9.1.4215 using a non-regenerative kinetics approach. Ligand (variant) injections were y-aligned and capture levels were determined by setting a report point after ligand capture. Curves were double-referenced, first by subtracting a negative control spot where no binding was expected to account for non-specific background interactions, followed by blanking with an injection of running buffer alone. A 1:1 Langmuir fit was applied to calculate the kd, ka, and kD of each variant.

#### C.9 Purification

For purification, remaining lysate (±900 µL) was mixed with 35 µL of MagneHis Ni-particles (Promega, V8565) in a 96-well plate (Thermo Scientific, AB1127). Purification was conducted using a Blue^®^Washer (BlueCatBio), which enables centrifugal removal of liquids from plates, including a magnetic plate to retain beads. After mixing the lysate with the beads, the lysate was spun off. Three cycles of washing were performed by adding 150 µL binding/wash buffer (100 mM HEPES, 10 mM imidazole, 500 mM NaCl), shaking for 2 minutes, and then removing the buffer while retaining the beads via magnetic-plate. After the third cycle, 50 µL of elution buffer (100 mM HEPES, 500 mM imidazole) were added to the wells, which were shaken again for 2 minutes before carefully pipetting the elution (while retaining the beads in the plate with a magnet) into clean 96-well plates.

#### C.10 Protein QC and normalization

Following purification, protein concentration was measured by bicinchoninic acid (BCA) assay (Merck, 71285-3) against a BSA standard curve for each well. Protein purity was determined by LabChip^®^ GXII Touch™, as above in the expression assay section, using 2 µL of purified protein. Peak analysis was performed using the LabChip GX Reviewer 5.14 software, and variants for which the protein of interest represented less than 55% of the overall protein in the purified protein were discarded. Based on the calculated concentration and percentage purity, proteins of interest were normalized in elution buffer to 400 ng/µL, though samples with concentrations between 200 ng/µL and 399 ng/µL were kept in the set given the high probability of still achieving a clear melting temperature measurement.

#### C.11 Thermostability assay

To measure melting temperature, the Prometheus NT.48 instrument (NanoTemper Technologies) was used. Standard capillaries (PR-C002, NanoTemper Technologies) were filled with approximately 10 µL of each purified protein sample and loaded into the sample drawer 48 at a time. Temperature was increased from 20°C to 80°C or 90°C by 1°C min-1. Melting temperatures were called manually based on the peak(s) observed from the first derivative of the ratio of the absorbance at 350 nm over the absorbance at 330 nm.

#### C.12 Serine-pyruvate aminotransferase activity assay

Enzymatic activity of serine-pyruvate aminotransferase was measured using a spectrophotometric assay detecting the formation of hydroxypyruvate via its irreversible reduction of water soluble tetrazolium (WST-1, Roche, 5015944001) into formazan, which absorbs at 480nm. Activity was measured in two separate reactions: one with a high substrate concentration, and another with a low substrate concentration. The high-concentration reaction was prepared with 800 nM of serine, and the low-concentration reaction with 100 nM of serine. Both reactions were also prepared with 25 nM of pyruvate, all in 100 mM Tris pH 7.5 buffer.

Prior to running the assay, 2.5 µL of normalised purified protein elutions were stamped into the reaction mix (prepared to 7.5 µL) for a total volume of 10 µL in 384-well clear flat-bottom assay plates, with the protein concentration at 100 ng/µL. Finally, 10 µL of WST-1 was added to all wells. Immediately, plates were placed in the Spark13 Multimode Microplate Reader (Tecan) and incubated for 90 minutes at 37°C, with absorbance measured every minute. All assays were run in triplicate. Reaction rates were calculated from the slope of the linear segment of the curve of absorbance over time.

#### C.13 Haloalkane dehalogenase activity assay

The enzymatic activity was measured using a fluorescence-based spectrophotometric assay as described in Dockalova, V. *et al*. ^44^. The assay monitors the increase in fluorescence intensity resulting from the cleavage of the halide ion from 4-bromomethyl-6,7-dimethoxycoumarin (COU-Br; Sigma-Aldrich, Cat. No. 301450) to yield the corresponding fluorescent product.

COU-Br was dissolved in DMSO to prepare a 1.25 mM working stock solution. Purified proteins were normalized to a final concentration of 0.16 µM and diluted in an enzyme mix containing 50 mM phosphate buffer (pH 7.4). For each reaction, 2 µL of 1.25 mM COU-Br stock solution was added to wells of a 384-well black microtiter plate (Corning, Cat. No. 3575). The reaction was initiated by adding 18 µL of the enzyme mix to each well, resulting in a final COU-Br concentration of 0.125 mM and a total reaction volume of 20 µL.

Fluorescence intensity was monitored continuously for 20 minutes at 30°C using a Spark^®^ Multimode Microplate Reader with excitation and emission wavelengths set to 345 nm and 437 nm, respectively (band-width 10 nm for both). Measurements were taken from the top of the plate. All assays were performed with two technical replicates and two biological replicates. Initial reaction rates were determined from the linear portion of the fluorescence curve within the first 200 seconds.

### D Software

As this work describes a multi-model workflow of a somewhat fiddly nature (including the automated training of models), we found that excellent software implementations were necessary to achieve the results described in this paper. For this reason we highlight several of the more important design principles that influenced our work:

**Monorepo of libraries:** our overall codebase is organized as a monorepo of Python libraries. Each library is independently versioned, describes its dependencies via a `pyprojeet.toml` file, provides a virtual environment specified via a `uv.loek` file (using `uv`^45^), and has its own unit tests via `pytest`^46^. Keeping each library within a single monorepo helps to encourage all team members to retain context across the whole codebase, whilst making cross-cutting changes easy to make. This is intentionally not ‘versioned at HEAD’ to make it possible to incrementally deploy upgrades or deprecate APIs across the codebase.

**Functional style:** a functional programming style is used throughout; mutation is almost entirely disallowed, and the ‘abstract/final pattern’^47^ is ubiquitously used to define interfaces (with enforcement via a custom extension of frozen dataclasses).

**Deterministic randomness:** all random numbers are generated using a splittable PRNG^48^, with a NumPy-based implementation providing an interface essentially identical to the random keys provided by JAX^49^. (Our choice of machine learning framework is PyTorch, largely for historical reasons.) This ensures near-perfect reproducibility (up to nondeterminism introduced via floating-point reductions) of all runs. This proved essential for debugging many edge cases.

**Buffer-offset-size protein representations:** batches of (potentially variable-length) protein sequences are ubiquitous in this system. These batches are represented as a triple (`buffer`, `offset`, `size`), where `buffer` is a 1-dimensional array of bytes (uint8s), and `offset` and `sizes` are equally-shaped n-dimensional arrays of int64s. This triple represents an n-dimensional ‘array of proteins’, for which the `i` th multi-index is considered to be the protein sequence stored at (in Python notation) `buffer[offset[i] offset[i] + size[i]]`. Each byte is interpreted as a ASCII-encoded character, which in turn is interpreted as one of the residues as per FASTA conventions, so that for example the byte `0×4d = 77 = M = methionine`. Additional tokens are treated in the same way, so that for example `0×23 = 35 = # = mask`. This offers a compact way to represent protein sequences that is both human-readable and machine-readable, for which model tokenization is zero-copy, and for which random-access is available via indexing (unlike if, for example, a mask token was represented as the multi-byte string `<mask>`, which is a common convention).

**Workflow orchestration:** the processes described in this paper require orchestrating a heterogeneous collection of models and other necessary computational tasks. Due to the ‘bursty’/’spiky’ nature of demand and the need for heterogeneous compute, we found traditional worker-queue architectures such as Celery50 to be a poor fit. Instead this is accomplished by using an in-house workflow orchestrator inspired by Flyte51, backed by a Kubernetes cluster running on Google Cloud.

**Code quality:** we use `ruff`^52^ to format and lint, `pyright`^53^ to typecheck, `ast-grep`^54^ to catch and ban known antipatterns, `pre-eommit`^55^ to run checks both locally and in CI, and `jaxtypin`^56^ to describe array shapes and dtypes.

We hope to provide individual technical blog posts describing some of these aspects in greater detail.

## E Glossary

Candidate sequence: variant of a template sequence; the output of the algorithm described in this paper
Diversify: synonym for ‘zero shot’
Function: synonym for ‘property’. Conventionally used in the expression ‘sequence-function data’, as opposed to ‘sequence-property data’
Language model: synonym for Logiter
Logiter: a model which consumes a masked sequence and provides a distribution over the 20 canonical amino acids at each masked position. ESM-2 would be a prototypical example
Masked sequence: a sequence for which some residues have been replaced with a mask token. Overall this is a word over an alphabet of size 21, consisting of the 20 amino acids and one mask token
Predictor: a model which consumes a sequence (without masks) and predicts one or more properties of that protein
Property: the result of an assay measuring a protein. Common examples are binding affinity, enzymatic activity, or expression
Residue: synonym for the 20 canonical amino acids. This term is used to distinguish these from non-physical sequence elements like mask tokens
Sequence: synonym for ‘protein’. In this work we mostly focus on single-chain proteins (without quaternary structure), making the term unambiguous
Target product profile: the collection of desired properties (at least this much binding, this much thermostability, …) for a lead optimization candidate (the ‘target product’) to be considered successful
Template sequence: the initial protein (or proteins), used as a starting point: this is modified to produce variants predicted to have improved function. Sometimes called the ‘lead’ or the ‘wild type’ in other sources
Zero shot: the first round of a lead optimization campaign, in which the only data available may be a template sequence. No wet lab measured data of protein properties is assumed to be available.

## Bibliography

1. Paul, S. M. et al. How to improve R&D productivity: the pharmaceutical industry’s grand challenge. Nature Reviews Drug Discovery 9, 203–214 (2010).

2. Zambaldi, V. et al. De novo design of highaffinity protein binders with AlphaProteo. arXiv (2024) doi:10.48550/arXiv.2409.08022.

3. Li, Q., Vlachos, E. N. & Bryant, P. Design of linear and cyclic peptide binders from protein sequence information. Communications Chemistry 8, 211 (2025).

4. Boyd, N., Guns, S. & Escalante Bio. Mosaic. https://github.com/escalantebio/mosaic (2025).

5. Stark, H. et al. BoltzGen: Toward Universal Binder Design. (2025).

6. Nabla Bio. JAM2: Fully computational design of druglike antibodies with high success rates. (2025).

7. Mille-Fragoso, L. S. et al. Efficient generation of epitope-targeted de novo antibodies with Germinal. bioRxiv (2025) doi:10.1101/2025.09.19.677421.

8. Chai Discovery Team et al. Zeroshot antibody design in a 24well plate. bioRxiv (2025) doi:10.1101/2025.07.05.663018.

9. Chai Discovery Team. Druglike antibody design against challenging targets with atomic precision. (2025).

10. Abramson, J. et al. Accurate structure prediction of biomolecular interactions with AlphaFold 3. Nature 630, 493–500 (2024).

11. Wohlwend, J. et al. Boltz1: Democratizing Biomolecular Interaction Modeling. bioRxiv (2024) doi:10.1101/2024.11.19.624167.

12. Passaro, S. et al. Boltz2: Towards Accurate and Efficient Binding Affinity Prediction. bioRxiv (2025) doi:10.1101/2025.06.14.659707.

13. Corso, G., Stärk, H., Jing, B., Barzilay, R. & Jaakkola, T. DiffDock: Diffusion Steps, Twists, and Turns for Molecular Docking. in International Conference on Learning Representations (ICLR) (2023).

14. Watson, J. L. et al. De novo design of protein structure and function with RFdiffusion. Nature 620, 1089–1100 (2023).

15. Dauparas, J. et al. Robust deep learning–based protein sequence design using ProteinMPNN. Science 378, 49–56 (2022).

16. Hie, B. L. et al. Efficient evolution of human antibodies from general protein language models. Nature Biotechnology 42, 275–283 (2024).

17. Funk, J. et al. ProteusAI: An OpenSource and UserFriendly Platform for Machine LearningGuided Protein Design and Engineering. bioRxiv (2024) doi:10.1101/2024.10.01.616114.

18. Jiang, K. et al. Rapid in silico directed evolution by a protein language model with EVOLVEpro. Science 387, (2025).

19. Zhang, Q. et al. Integrating protein language models and automatic biofoundry for enhanced protein evolution. Nature Communications 16, 1553 (2025).

20. Frey, N. C. et al. Labintheloop therapeutic antibody design with deep learning. bioRxiv (2025) doi:10.1101/2025.02.19.639050.

21. Lin, Z. et al. Evolutionaryscale prediction of atomiclevel protein structure with a language model. Science 379, 1123–1130 (2023).

22. Sasaki, T., Hiroki, K. & Yamashita, Y. The Role of Epidermal Growth Factor Receptor in Cancer Metastasis and Microenvironment. BioMed Research International 2013, (2013).

23. Cotet, T.S. et al. Crowdsourced Protein Design: Lessons From the Adaptyv EGFR Binder Competition. bioRxiv (2025) doi:10.1101/2025.04.17.648362.

24. Hanke, L. et al. An alpaca nanobody neutralizes SARSCoV2 by blocking receptor interaction. Nature Communications 11, 4420 (2020).

25. Chippaux, J.P. Estimate of the burden of snakebites in subSaharan Africa: A metaanalytic approach. Toxicon 57, 586–599 (2011).

26. Suraweera, W. et al. Trends in snakebite deaths in India from 2000 to 2019 in a nationally representative mortality study. eLife 9, (2020).

27. Ahmadi, S. et al. Nanobody-based recombinant antivenom for cobra, mamba and rinkhals bites. Nature 647, 716–725 (2025).

28. Funk, J. et al. ProteusAI: An OpenSource and UserFriendly Platform for Machine LearningGuided Protein Design and Engineering. bioRxiv (2024) doi:10.1101/2024.10.01.616114.

29. Suzek, B. E., Huang, H., McGarvey, P., Mazumder, R. & Wu, C. H. UniRef: comprehensive and non redundant UniProt reference clusters. Bioinformatics 23, 1282–1288 (2007).

30. Biswas, S., Khimulya, G., Alley, E. C., Esvelt, K. M. & Church, G. M. LowN protein engineering with data efficient deep learning. Nature Methods 18, 389–396 (2021).

31. Ferragu, C. et al. gDPO: Scalable Preference Optimization for Protein Language Models. arXiv (2025) doi:10.48550/arXiv.2510.19474.

32. Jia, L., Jain, M. & Sun, Y. Improving antibody thermostability based on statistical analysis of sequence and structural consensus data. Antibody Therapeutics 5, 202–210 (2022).

33. Uçar, T. & Sormanni, P. BLOSUM Is All You Learn — Generative Antibody Models Reflect Evolutionary Priors. bioRxiv (2025) doi:10.1101/2025.10.26.684652.

34. Uçar, T., Malherbe, C. & Gonzalez, F. Exploring LogLikelihood Scores for Ranking Antibody Sequence Designs. bioRxiv (2025) doi:10.1101/2024.10.07.617023.

35. Wei, J. et al. Emergent Abilities of Large Language Models. Transactions on Machine Learning Research (2022).

36. Parnami, A. & Lee, M. Learning from Few Examples: A Summary of Approaches to FewShot Learning. arXiv doi:10.48550/arXiv.2203.04291.

37. Faw, M., Sen, R., Zhou, Y. & Das, A. InContext FineTuning for TimeSeries Foundation Models. in Forty-second International Conference on Machine Learning (2025).

38. Cradle EGFR Competition Follow Up. https://proteinbase.com/collections/cradleegfrcompetition (2025).

39. Jumper, J. et al. Highly accurate protein structure prediction with AlphaFold. Nature 596, 583–589 (2021).

40. Richardson, L. et al. MGnify: the microbiome sequence data analysis resource in 2023. Nucleic Acids Research 51, D753–D759 (2022).

41. Gao, S. H., Huang, K., Tu, H. & Adler, A. S. Monoclonal antibody humanness score and its applications. BMC Biotechnology 13, 55 (2013).

42. Jensen, K. K. et al. Improved methods for predicting peptide binding affinity to MHC class II molecules. Immunology 154, 394–406 (2018).

43. Katoh, K., Misawa, K., Kuma, K. & Miyata, T. MAFFT: a novel method for rapid multiple sequence alignment based on fast Fourier transform. Nucleic Acids Research 30, 3059–3066 (2002).

44. Dockalova, V. et al. Fluorescent substrates for haloalkane dehalogenases: Novel probes for mechanistic studies and protein labeling. Computational and Structural Biotechnology Journal 18, 922–932 (2020).

45. Astral.sh. uv. https://github.com/astralsh/uv/.

46. Krekel, H. et al. pytest. https://github.com/pytestdev/pytest (2004).

47. Kidger, P. The abstract/final pattern. https://docs.kidger.site/equinox/pattern/ (2024).

48. Salmon, J. K., Moraes, M. A., Dror, R. O. & Shaw, D. E. Parallel random numbers: as easy as 1, 2, 3. in Proceedings of 2011 International Conference for High Performance Computing, Networking, Storage and Analysis (Association for Computing Machinery, Seattle, Washington, 2011). doi:10.1145/2063384.2063405.

49. Bradbury, J. et al. JAX: composable transformations of Python+NumPy programs. http://github.com/jaxml/jax (2018).

50. Celery. https://github.com/celery/celery.

51. Union.ai. Flyte. https://flyte.org/.

52. Astral.sh. ruff. https://github.com/astralsh/ruff/.

53. Microsoft. pyright. https://github.com/microsoft/pyright/.

54. astgrep. https://ast-grep.github.io/.

55. precommit. https://precommit.com/.

56. Kidger, P. jaxtyping. http://github.com/patrickkidger/jaxtyping.

